# Vesicular and non-vesicular extracellular small RNAs direct gene silencing in a plant-interacting bacterium

**DOI:** 10.1101/863902

**Authors:** Antinéa Ravet, Jérôme Zervudacki, Meenu Singla-Rastogi, Magali Charvin, Odon Thiebeauld, Alvaro L Perez-Quintero, Lucas Courgeon, Adrien Candat, Liam Lebeau, Antonio Emidio Fortunato, Venugopal Mendu, Lionel Navarro

**Author notes:** These authors contributed equally to the work (Antinéa Ravet, Jérôme Zervudacki).

## Abstract

Extracellular plant small RNAs (sRNAs) and/or double-stranded RNA (dsRNA) precursors act as triggers of RNAi in interacting filamentous pathogens^1–7^. However, whether any of these extracellular RNA species direct gene silencing in plant-interacting bacteria remains unknown. Here, we show that Arabidopsis transgenic plants expressing sRNAs directed against virulence factors of a *Pseudomonas syringae* strain, reduce its pathogenesis. This Antibacterial Gene Silencing (AGS) phenomenon is directed by Dicer-Like (DCL)-dependent antibacterial sRNAs, but not cognate dsRNA precursors. Three populations of active extracellular sRNAs were recovered in the apoplast of these transgenic plants. The first one is mainly non-vesicular and associated with proteins, whereas the second one is likely located inside Extracellular Vesicles (EVs). Intriguingly, the third population is unbound to proteins and in a dsRNA form, unraveling a novel class of functional extracellular free sRNAs (efsRNAs). Both Arabidopsis transgene- and genome-derived efsRNAs were retrieved inside bacterial cells. Finally, we show that salicylic acid (SA) promotes AGS, and that a substantial set of endogenous efsRNAs exhibits predicted bacterial targets that are down-regulated by SA biogenesis and/or signaling during infection. This study thus unveils an unexpected AGS phenomenon, which may have wider implications in the understanding of how plants regulate transcriptome, microbial community composition and genome evolution of associated bacteria.

## Introduction

RNAi is a conserved gene regulatory mechanism that promotes antiviral defense by repressing translation, accumulation and replication of viral RNAs^8^. In plants, RNAi also orchestrates resistance against bacterial, fungal and oomycetal pathogens, partly by fine-tuning the expression of immune-responsive genes^9,10^. The core mechanism of RNAi involves the processing of double-stranded RNAs (dsRNAs) by DCL proteins, leading to the production of 20-25 nt long small RNAs (sRNAs). Small RNAs are then loaded into Argonaute (AGO) proteins to direct post-transcriptional silencing of sequence-complementary mRNA targets, through endonucleolytic cleavage and/or translational inhibition^11^.

An important feature of plant sRNAs is their ability to trigger non-cell autonomous silencing in adjacent cells as well as in distal tissues^12,13^. This phenomenon is essential to prime antiviral response ahead of the infection front, but also to translocate silencing signals between plant cells and their non-viral eukaryotic interacting (micro)organisms^1,14^. For example, plant sRNAs are exported in the fungal pathogens *Verticillium dahliae*^2^ and *Botrytis cinerea*^3^, as well as in the oomycetal pathogen *Phytophthora capsici*^4^, to silence pathogenicity factors. The transfer of sRNAs from plant cells towards *B. cinerea* and *P. capsici* cells is in part mediated by extracellular vesicles (EVs)^1,4,5^. Conversely, fungal sRNAs from *B. cinerea*, and oomycetal sRNAs from *Hyaloperonospora arabidopsidis*, are translocated into plant cells to silence defense genes^15,16^. In addition, some transfer RNA-derived fragments (tRFs), produced by the rhizobium *Bradyrhizobium japonicum*, can silence genes in the soybean *Glycine max* to promote nodulation^17^. However, whether plant sRNAs could be transferred in interacting bacteria, and in turn directly regulate bacterial gene expression, remains elusive.

Artificial trans-kingdom RNAi has long been employed to direct Host-Induced Gene Silencing (HIGS), a technology used to characterize the function of virulence genes or to engineer disease resistance in plants^3^. HIGS notably relies on *in planta* expression of dsRNAs bearing homologies to essential and/or virulence genes, and can operate in insects, nematodes, parasitic plants, oomycetes, and fungi^18^. For example, HIGS confers full protection against *Fusarium graminearum*^6^ and *B. cinerea*^3^, a phenotype that can be recapitulated by spraying antifungal dsRNAs and/or sRNAs onto wild-type plants prior to infection. The latter Spray-Induced Gene Silencing (SIGS) phenomenon is reminiscent of environmental RNAi, a process that involves the uptake of RNAs from the environment to trigger RNAi. However, so far, HIGS and SIGS have been shown to be effective in eukaryotic (micro)organisms possessing canonical RNAi factors. Indeed, there is no evidence indicating that the stable *in planta* expression, or the external application, of artificial sRNAs and/or dsRNAs could trigger gene silencing in a plant-interacting bacterium.

Here, we used the Gram-negative pathogenic bacterium *Pseudomonas syringae* pv. *tomato* strain DC3000 (*Pto* DC3000), and the plant *Arabidopsis thaliana*, as a model interaction system to address these questions. We show that the stable *in planta* expression of an inverted repeat transgene generating sRNAs directed against two *Pto* DC3000 virulence factors can dampen bacterial pathogenesis. Three populations of active extracellular sRNAs were recovered from the apoplastic fluid. The first one is mostly associated with proteins that are located outside PENETRATION1 (PEN1)-positive EVs, while the second one is likely located inside TETRASPANIN8 (TET8)-positive EVs. Intriguingly, the third one is unbound to proteins and exists in a dsRNA form. Importantly, a subset of these artificial extracellular free sRNAs (efsRNAs) was found internalized by *Pto* DC3000 cells. This was also true for endogenous efsRNAs produced from various Arabidopsis genomic origins. Finally, we show that SA promotes AGS activity, and that a substantial set of internalized endogenous sRNAs exhibits sequence-complementarity to *Pto* DC3000 mRNAs, whose cognate protein levels were found down-regulated by SA biogenesis and/or signaling during infection. Overall, this work unveils the vesicular and non-vesicular extracellular sRNAs that are causal for AGS. It also highlights a potential for endogenous plant sRNAs in directly reprogramming gene expression in plant-associated bacteria.

## Results

### Stable expression of anti-*cfa6*/*hrpL* sRNAs in Arabidopsis reduces *Pto* DC3000 virulence

*Pto* DC3000 is the causal agent of bacterial speck in tomato and can also infect *Arabidopsis thaliana*^18^. This bacterium enters Arabidopsis leaf tissues through stomata, and subsequently reaches the apoplast, where it multiplies at high population levels^19^. Upon detection of *Pto* DC3000, Arabidopsis triggers stomatal closure within an hour of infection, which limits the access of this bacterium into inner leaf tissues^20,21^. As a counter-defense, *Pto* DC3000 actively reopens stomata at 3 hours post-inoculation (hpi), in part, by secreting the phytotoxin coronatine (COR)^19,22^. Accordingly, a *Pto* DC3000 Δ*cfa6* (*Pto*Δ*cfa6*) strain that is deleted in *Cfa6*, a gene encoding a structural component of COR^23^, does not trigger stomatal reopening on wild-type Columbia-0 (Col-0) leaf sections at 3 hpi (Fig. 1a). A similar phenotype is observed in response to a *Pto* DC3000 Δ*hrpL* (*Pto*Δ*hrpL*) strain that is deleted in *HrpL* (Fig. 1b). The latter gene encodes an alternative sigma factor that directly controls the expression of type III secretion system (TTSS)-associated genes, and indirectly the expression of *COR* biosynthesis genes^24,25^. Both phenotypes are rescued upon exogenous application of COR (Fig. 1a, b), indicating that they are caused by the inability of these mutant strains to produce COR^24,25^. Conversely, and as shown previously^20^, a normal stomatal reopening phenotype occurs during infection with a type III secretion-defective *Pto* DC3000 Δ*hrcC* mutant (*Pto*Δ*hrcC*) (Fig. 1b).

**Figure 1.**
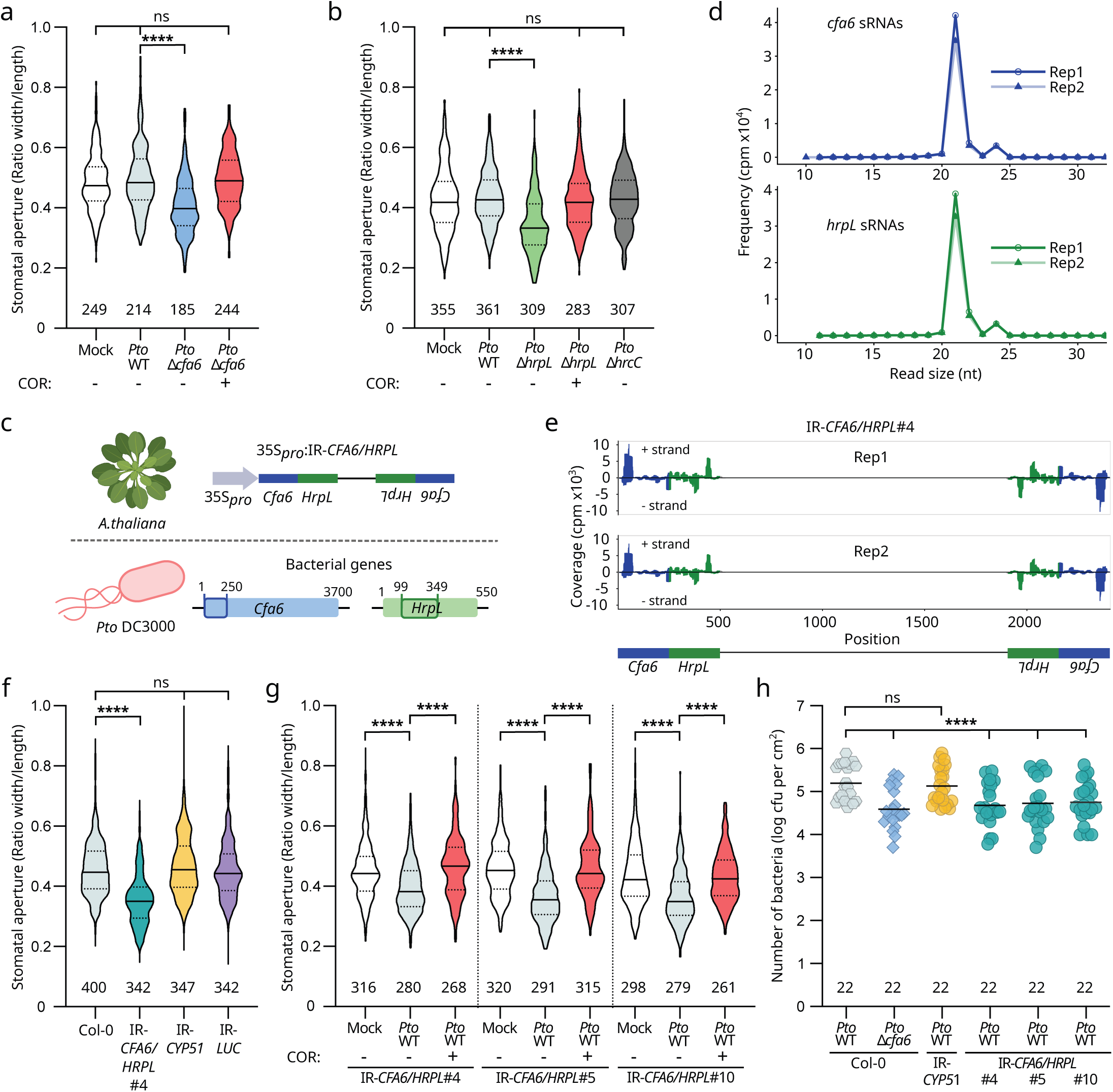
Stable expression of the IR-*CFA6/HRPL* transgene in Arabidopsis reduces *Pto* DC3000 virulence during infection. **a.** The coronatine (COR)-defective *Pto*Δ*cfa6* strain is impaired in its ability to reopen stomata and this phenotype is rescued upon exogenous application of COR. Leaf sections of Col-0 plants were treated with mock (water), WT *Pto* DC3000 (*Pto* WT) or *Pto*Δ*cfa6* –with or without COR– and the stomatal aperture response was analyzed at 3 hours post-inoculation (hpi). The concentrations of bacterial strains and of COR were of 10^8^ cfu mL^-1^ and 1μM, respectively. The number of stomata analyzed per condition is written underneath each condition and statistical significance was assessed using a one-way ANOVA test (ns: p-value ≥ 0.05; ****: p-value < 0.0001). Similar results were obtained in three independent experiments and are pooled in a single plot. **b.** The *Pto*Δ*hrpL* strain, but not the *Pto*Δ*hrcC* strain, is impaired in its ability to reopen stomata and this phenotype is rescued upon exogenous application of COR. The stomatal aperture assay was conducted as in a. with leaf sections of Col-0 inoculated with *Pto* WT, Δ*hrpL* –with or without COR– and Δ*hrcC* strains. **c.** Schematic representations of the IR-*CFA6*/*HRPL* chimeric hairpin construct stably expressed in Arabidopsis transgenic plants (upper panel), and of the 250 bp regions of the *Cfa6* (1-250 nt, blue box) and *HrpL* (99-349 nt, green box) genes that are targeted by sRNAs in *Pto* DC3000 (bottom panel). **d.** Size distribution and abundance of anti-*cfa6*/*hrpL* sRNA reads along the *CFA6/HRPL* inverted repeat, sequenced from the IR-*CFA6*/*HRPL*#4 plants. **e.** Total sRNA reads that map to the IR-*CFA6*/*HRPL* inverted repeat are depicted. Anti-*cfa6* and anti-*hrpL* sRNAs are shown in blue and green, respectively. **f.** *Pto* DC3000 WT no longer induces stomatal reopening in Arabidopsis IR-*CFA6*/*HRPL* transgenic lines. Stomatal aperture measurements were conducted on leaf sections of IR-*CFA6/HRPL*#4, IR-*CYP51* and IR-*LUC* lines infected with *Pto* WT, as described in a. **g.** The impaired stomatal reopening phenotypes observed in IR-*CFA6*/*HRPL* lines are rescued upon application of exogenous COR. Stomatal aperture measurements were conducted on leaf sections of IR-*CFA6/HRPL*#4, #5, #10 lines infected with *Pto* WT, as described in a., with or without COR. **h.** Arabidopsis IR-*CFA6/HRPL* lines exhibit reduced *Pto* WT titers when compared to Col-0 and IR*-CYP51-*infected plants at 2 days post-infection (dpi). Five to six-week-old Col-0, IR*-CYP51* and IR-*CFA6/HRPL*#4, #5 and #10 plants were dip-inoculated, at a concentration of 5 x 10^7^ cfu mL^-1^, with *Pto* WT-GFP. As a control condition, Col-0 plants were treated at the same concentration with *Pto*Δ*cfa6*-GFP. Number of leaf disc analyzed is written underneath each condition. Data from three independent experiments are pooled in a single plot. Statistical significance was assessed using a two-way ANOVA test (ns: p-value>0.05; ****: p-value<0.0001).

Because both *Cfa6* and *HrpL* genes are critical for *Pto* DC3000 pathogenesis, and only expressed during infection^23–25^, we reasoned that they represent suitable targets to test the possible occurrence and relevance of AGS. We thus decided to stably express anti-*cfa6* and anti-*hrpL* sRNAs from Arabidopsis and first monitored their possible anti-virulence effects through a stomatal reopening assay. More specifically, we generated Arabidopsis transgenic lines expressing a chimeric inverted repeat, bearing homologies with the coding regions of both *Cfa6* and *HrpL* genes (Fig. 1c). As a negative control, we generated Arabidopsis transgenic lines expressing inverted repeats targeting three cytochrome P450 lanosterol C-14α-demethylase (*CYP51*) genes of the fungus *F. graminearum*^6,7^, or the luciferase (*LUC*) reporter gene. All the engineered inverted repeat transgenes were driven by the Cauliflower Mosaic Virus (CaMV) 35S promoter. The resulting stable transgenic lines, respectively referred to as IR-*CFA6/HRPL*, IR-*CYP51* or IR-*LUC*, expressed the cognate engineered artificial sRNAs, and did not exhibit developmental defects compared to Col-0 plants (Supplementary Fig. 1a-c). Small RNA sequencing (sRNA-seq) from the IR-*CFA6/HRPL*#4 reference line revealed typical peaks of 21 and 24 nt endogenous sRNAs mapping to the Col-0 genome (Supplementary Fig. 1d), and a high accumulation of 21 nt anti-*cfa6*/*hrpL* sRNAs (Fig. 1d), which were produced along the *CFA6*/*HRPL* regions of the hairpin (Fig. 1e). Further target prediction analysis of anti-*cfa6*/*hrpL* sRNAs against the Col-0 and *Pto* DC3000 annotated genes, coupled with the analysis of free energy of corresponding sRNA-target pairings, indicated that off-target effects are unlikely (Supplementary Fig. 1e, Supplementary Table 1 and 2).

We further monitored *Pto* DC3000-induced stomatal reopening response in IR-*CFA6/HRPL* transgenic lines. Importantly, these transgenic lines were found insensitive to the stomatal reopening response triggered by *Pto* DC3000 at 3 hpi (Fig. 1f), thereby mimicking the phenotypes observed during infection of Col-0 with the *Pto*Δ*cfa6* or *Pto*Δ*hrpL* strains (Fig. 1a, b). By contrast, the *Pto* DC3000-induced stomatal reopening phenotype was unaltered in the infected IR-*CYP51* and IR-*LUC* lines, compared to Col-0-infected plants (Fig. 1f), supporting a sequence-specific effect. Furthermore, the compromised stomatal reopening phenotypes detected in three independent IR-*CFA6/HRPL*-infected lines were rescued upon exogenous application of COR (Fig. 1g). Therefore, these phenotypes are presumably caused by an altered ability of the leaf-associated and/or surrounding *Pto* DC3000 cells to produce COR.

We next reasoned that, by suppressing stomatal reopening, anti-*cfa6*/*hrpL* sRNAs would limit the entry of *Pto* DC3000 into leaf tissues, and consequently reduce its apoplastic colonization. To test this, we dip-inoculated Col-0, IR-*CYP51* and three independent IR-*CFA6/HRPL* lines with *Pto* DC3000 and monitored bacterial titers in those plants. It is noteworthy that *Pto* DC3000 infection did not alter the accumulation of anti-*cfa6*/*hrpL* in these conditions (Supplementary Fig. 1f). We found that *Pto* DC3000 was less effective in colonizing the apoplast of IR-*CFA6/HRPL* lines compared to Col-0- and IR-*CYP51*-infected plants, thereby mimicking the growth defect of *Pto*Δ*cfa6* on Col-0 plants (Fig. 1h). Altogether, these data indicate that the expression of sRNAs directed against *Cfa6* and *HrpL* genes in Arabidopsis plants can reduce the virulence functions of *Pto* DC3000 during infection.

### Antibacterial sRNAs, but not cognate dsRNA precursors, are biologically active

To get a first insight into the active RNA entities responsible for AGS, we introduced the *dcl2*-*1*, *dcl3*-*1* and *dcl4*-*2* mutations in a reference line expressing an IR-*HRPL* inverted repeat under the control of the constitutive Ubi.U4 promoter^26^. It is noteworthy that this promoter was used here, instead of the 35S promoter, to prevent co-suppression effect caused by the presence of 35S copies in the T-DNA insertional mutant lines^27,28^. A drastic reduction in the accumulation of anti-*hrpL* sRNAs was observed in Ubi.U4::IR-*HRPL dcl234* compared to Ubi.U4::IR-*HRPL* control plants, as revealed by northern blot analysis (Fig. 2a, left panel). Furthermore, unprocessed forms of IR-*HRPL* precursors accumulated in Ubi.U4::IR-*HRPL dcl234* plants, while they were barely detectable in the Ubi.U4::IR-*HRPL* parental line (Fig. 2a, right panel). Collectively, these data support a DCL-dependent processing of *HRPL* dsRNAs into anti-*hrpL* siRNAs. Importantly, when the Ubi.U4::IR-*HRPL dcl234* plants were used in a stomatal reopening assay, we found that *Pto* DC3000 triggered a normal stomatal reopening phenotype, similar to the effect observed in Col-0 and *dcl234* control plants (Fig. 2b). By contrast, a full suppression of stomatal reopening events was achieved in the Ubi.U4::IR-*HRPL* reference line (Fig. 2b), which produces abundant anti-*hrpL* sRNAs (Fig. 2a). Collectively, these data provided evidence that long *HRPL* dsRNAs are unlikely responsible for the anti-virulence effect, and rather suggested that sRNA species are the RNA entities responsible for AGS. To further test this assumption, we decided to establish a semi-*in vitro* assay. Col-0 leaf sections were pre-treated with IR-*CFA6*/*HRPL*#4 total RNA extracts for one hour and were subsequently challenged with *Pto* DC3000, following which the stomatal aperture events were monitored at 3 hpi. A full suppression of the *Pto* DC3000-induced stomatal aperture was detected in response to these RNA extracts (Fig. 2c). However, the stomatal reopening phenotype remained unaltered in response to RNA extracts derived from Col-0, IR-*CYP51* and IR-*LUC* plants, supporting sequence-specificity (Fig. 2c). Therefore, the pretreatment of IR-*CFA6*/*HRPL*#4 RNA extracts on Col-0 leaf sections recapitulates the anti-virulence effect achieved by IR-*CFA6*/*HRPL*#4 leaf sections (Fig. 1f, g). We further conducted size separation of the IR-*CFA6*/*HRPL*#4 total RNA extracts and tested the activities of the resulting RNA fractions. The long RNA fraction (>200 nt), which presumably contains long *CFA6*/*HRPL* dsRNAs, did not suppress *Pto* DC3000-triggered stomatal reopening (Fig. 2d; Supplementary Fig. 2a, b). In contrast, the small RNA fraction (<200 nt), likely containing anti-*cfa6*/*hrpL* sRNAs, was as active as IR-*CFA6*/*HRPL*#4 total RNA extracts (Fig. 2d). Collectively, these data indicate that anti-*cfa6*/*hrpL* sRNAs must be the active RNA entities from the IR-*CFA6*/*HRPL*#4 total RNA extracts. To confirm this, we made use of the viral protein p19, whose property is to selectively bind sRNA duplexes^29,30^. Chitin magnetic beads coated with p19 proteins were incubated with total RNA extracts from IR-*LUC* and IR-*CFA6/HRPL*#4 plants. The eluted bound sRNA duplexes were subsequently analyzed by northern blot. Large amounts of pulled-down anti*-cfa6/hrpL* sRNAs, and anti*-luc* control sRNAs, were detected through this approach (Fig. 2e). Bound sRNA species were mostly 21 nt in length, which is consistent with the high affinity of p19 for 21 nt sRNA duplexes^29^. Importantly, pulled-down anti-*cfa6*/*hrpL* sRNAs were fully capable of suppressing stomatal reopening, such as IR-*CFA6*/*HRPL*#4 total RNA extracts (Fig. 2f). In contrast, the pulled-down anti-*luc* sRNAs were found inactive (Fig. 2f). These data further support that anti-*cfa6*/*hrpL* sRNA duplexes, but not anti-*luc* sRNAs, harbour anti-virulence activity against *Pto* DC3000. We finally synthesized *CFA6*/*HRPL* dsRNAs and anti-*cfa6*/*hrpL* sRNA duplexes *in vitro* (Fig. 2g), and tested their activities. *In vitro*-synthesized *CFA6/HRPL* dsRNAs did not alter the ability of *Pto* DC3000 to reopen stomata, nor did *CYP51* control dsRNAs (Fig. 2h). In contrast, *in vitro*-synthesized anti-*cfa6/hrpL* sRNAs suppressed stomatal reopening events, which was not the case with anti-*cyp51* sRNAs (Fig. 2h). Altogether, these data provide solid evidence that sRNAs, but not their unprocessed dsRNA precursors, are biologically active.

**Figure 2.**
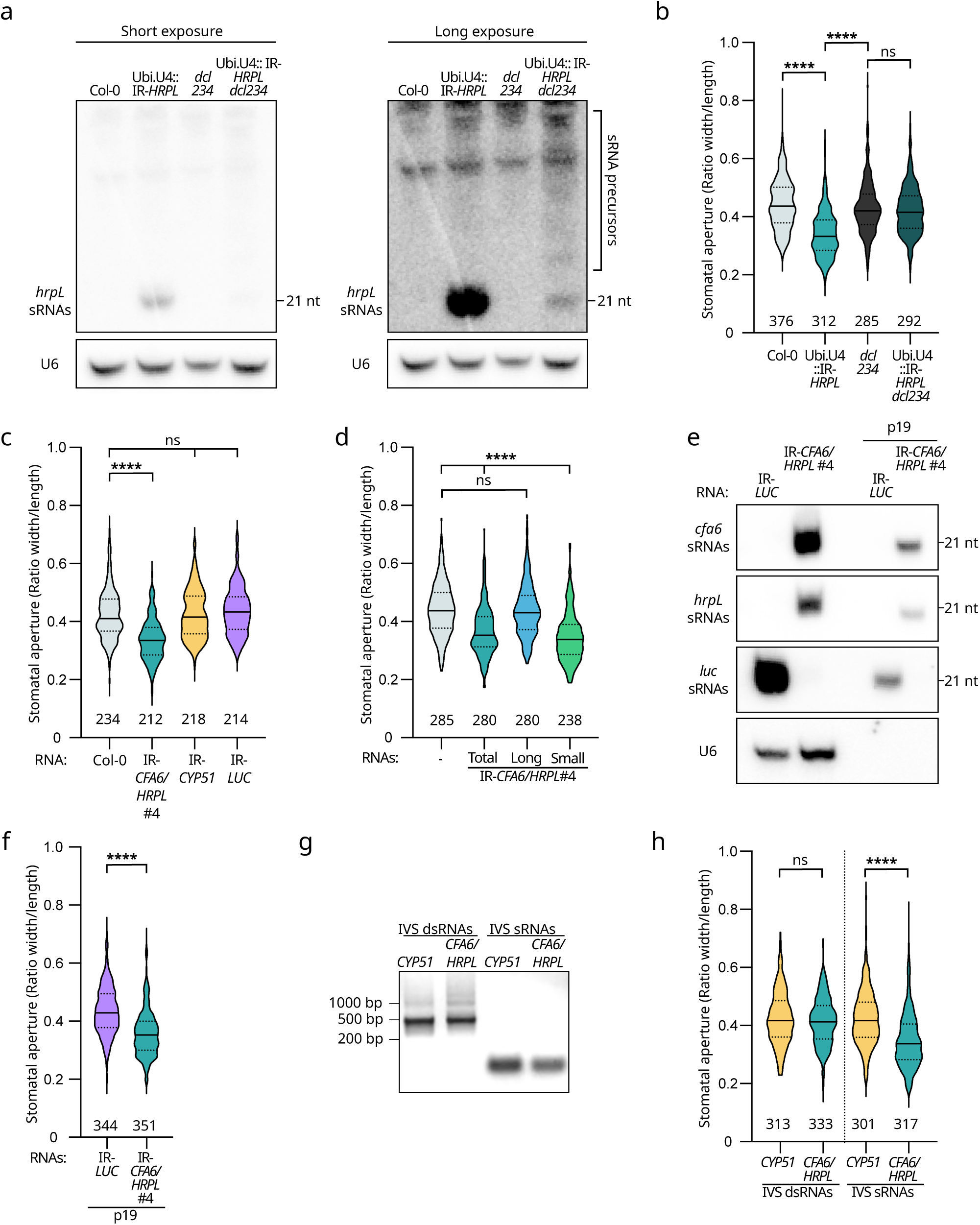
Antibacterial sRNA species, but not unprocessed dsRNAs, are causal for the suppression of *Pto* DC3000-triggered stomatal aperture. **a**. Accumulation level of anti-*hrpL* sRNAs in Col-0, Ubi.U4 IR-*HRPL*, *dcl2-1 dcl3-1 dcl4-2* (*dcl234*), and Ubi.U4 IR-*HRPL* in *dcl234* mutant background (Ubi.U4 IR-*HRPL dcl234*) was assessed by low molecular weight northern blot analysis. The northern blot results, obtained from the same membrane, are shown at two different expositions: short (left) and long (right). The long exposition was depicted to better visualize unprocessed forms of dsRNA precursors, referred to as “sRNA precursors” in the panel. Because the same membrane was used to obtain the results shown in panel a, the same U6 loading control This is one representative experiment out of the three independent experiments performed. **b.** Ubi.U4 IR-*HRPL dcl234* plants do not suppress *Pto* DC3000-induced stomatal reopening response. The stomatal reopening assay was conducted on leaf sections of the same genotypes detailed in a. and as described in Fig 1a. Similar results were obtained in three independent experiments and are pooled in a single plot. The number of stomata analyzed per condition is written underneath each condition and statistical significance was assessed using a one-way ANOVA test (ns: p-value ≥ 0.05; ****: p-value < 0.0001). **c.** Twenty ng μl^-1^ of total RNA from IR-*CFA6/HRPL#4* but not IR-*LUC* and IR-*CYP51* prevents stomatal reopening. Col-0 leaf sections were incubated with total RNA from Col-0, IR-*CFA6/HRPL#4,* IR-*LUC* or IR-*CYP51* and stomatal reopening assay was performed as in b. Similar results were obtained in two independent experiments and are pooled in a single plot. **d.** Small RNA species, but not corresponding long RNA species, from IR-*CFA6/HRPL*#4 plants suppress *Pto* DC3000-induced stomatal reopening. The stomatal reopening assay was conducted with *Pto* DC3000 and by incubating Col-0 leaf sections with total, long (> 200 nt) or small (< 200 nt) RNA fractions, separated from total RNAs of IR-*CFA6/HRPL*#4 plants. Similar results were obtained in three independent experiments and are pooled in a single plot. **e.** Accumulation of anti-*cfa6*/*hrpL* sRNAs in IR-*CFA6/HRPL*#4 and IR-*LUC* total RNAs before or after p19 pull-down was detected by low molecular weight northern blot analysis. U6 was used as a control of intracellular RNAs. This is one representative experiment out of the three independent experiments performed. **f.** P19 pulled-down sRNAs from the IR-*CFA6/HRPL*#4 total RNAs suppress the ability of *Pto* DC3000 to reopen stomata. Stomatal aperture assay was conducted as in b. Statistical significance was assessed using Student’s *t*-test (****: p-value<0.0001). Similar results were obtained in three independent experiments and are pooled in a single plot. **g.** Agarose gel picture of ethidium bromide-stained *in vitro* synthesized (IVS) long dsRNAs and sRNA duplexes. **h.** *In vitro* synthesized anti-*cfa6*/*hrpL* sRNAs, but not cognate unprocessed dsRNA precursors, suppress *Pto* DC3000-triggered stomatal reopening. The stomatal reopening assay was conducted as in b. Similar results were obtained in three independent experiments and are pooled in a single plot.

### Anti-*hrpL* sRNAs are causal for the suppression of stomatal reopening

Although the above findings indicate that anti-*cfa6*/*hrpL* sRNAs are effective against *Pto* DC3000, they do not firmly demonstrate that they are causal for this phenomenon. To address this, we expressed, in the *Pto*Δ*hrpL* mutant, either a WT *HrpL* transgene or a mutated (mut *HrpL*) version that contains as many silent mutations as possible in the sRNA targeted region (Fig. 3a, b). These mutations are expected to alter the binding of anti-*hrpL* sRNAs, as revealed by just a few predicted sRNA-mut *HrpL* interactions, whose free energy mean was higher compared to the one of the numerous sRNA-*HrpL* interactions (Fig. 3c, Supplementary Table 3). Both transgenes were expressed under the constitutive neomycin phosphotransferase II (*NPTII*) promoter (Fig. 3a). The resulting recombinant bacteria, referred to here as *Pto*Δ*hrpL* WT *HrpL* and *Pto*Δ*hrpL* mut *HrpL*, restored the ability of *Pto*Δ*hrpL* strain to trigger the reopening of stomata (Fig. 3d), indicating that both transgenes are functional in a stomatal reopening assay. To assess the specific effect of sRNAs towards suppression of *HrpL*-mediated stomatal aperture, we generated two independent Arabidopsis 35S::IR-*HRPL* transgenic lines, referred to as IR-*HRPL*#1 and #4. These lines overexpressed anti-*hrpL* sRNAs (Supplementary Fig. 3), and, expectedly, suppressed the ability of *Pto* DC3000 to reopen stomata (Fig. 3e). Similar results were obtained in response to *Pto*Δ*hrpL* WT *HrpL* (Fig. 3e), supporting a sensitivity of this bacterial strain to sRNA action. By contrast, the *Pto*Δ*hrpL* mut *HrpL* strain was fully competent in reopening the stomata (Fig. 3e), indicating that anti-*hrpL* sRNAs no longer exert their anti-virulence effects towards this bacterium. Altogether, these data demonstrate that anti-*hrpL* sRNAs are causal for the suppression of *HrpL*-mediated stomatal reopening function.

**Figure 3.**
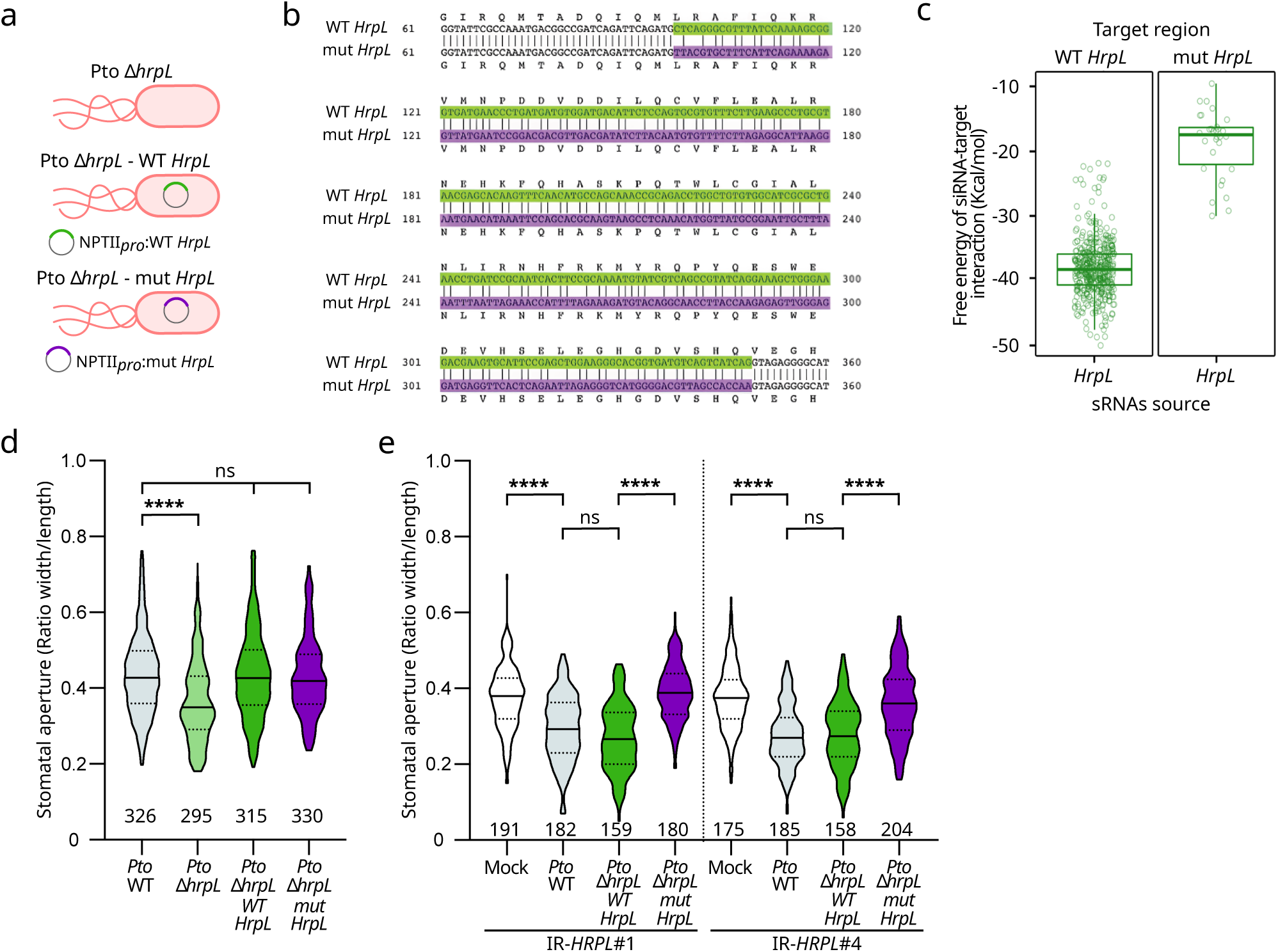
Anti-*hrpL* sRNAs are causal for the suppression of stomatal reopening. **a.** Schematic representation of *Pto*Δ*hrpL* and of the recombinant strains expressing either the WT *HrpL* or mut *HrpL* constructs, under the control of the constitutive *NPTII* promoter. **b.** The sequence of WT *HrpL* (99-349 nt) selected to generate the inverted repeat transgene (highlighted in green) was aligned with the sequence of mutated *HrpL* (mut *HrpL*) designed to contain as many silent mutations as possible in the sRNA targeted region (highlighted in purple). **c.** Thermodynamic energy analysis of sRNA-target interactions between anti-*hrpL* sRNAs and the WT or mutated *HrpL* sequence versions. **d.** Both the WT *HrpL* and mut *HrpL* constructs fully complemented *PtoΔhrpL* for its ability to reopen stomata. Col-0 leaves were incubated with *PtoΔhrpL*, *PtoΔhrpL* WT *HrpL* and *PtoΔhrpL* mut *HrpL* bacterial strains. Stomatal reopening response was assessed as described previously. Statistical significance was assessed using a one-way ANOVA test (ns: p-value ≥ 0.05; ****: p-value < 0.0001). Similar results were obtained from three independent experiments and the data are pooled in a single plot. **e.** The *Pto*Δ*hrpL* mut *HrpL* strain is refractory to anti-*hrpL* sRNAs action at 3 hpi. The stomatal reopening assay was conducted as in d., IR-*HRPL*#1 and #4 leaf sections were treated with mock (water) or the indicated bacteria at a concentration of 10^8^ cfu mL^-1^. Results of two independent experiments are pooled in a single plot. Statistical significance was assessed using a one-way ANOVA test (ns: p-value>0.05; ****: p-value<0.0001).

### Active apoplastic particles that co-purify with PEN1-positive EVs are likely composed of protein-bound anti-*cfa6*/*hrpL* sRNAs located outside these EVs

Since apoplastic EVs were previously shown to contribute to the trafficking of plant sRNAs towards fungal and oomycetal pathogens^1,4^, we investigated whether they could similarly participate in the transfer of active sRNAs from Arabidopsis towards *Pto* DC3000. Two main categories of plant EVs have been characterized so far. The first one is composed of vesicles that are heterogenous in size, and that are notably marked by PEN1^31^. The second one, which is required for the transfer of active sRNAs from Arabidopsis towards *B*. *cinerea*^1,5^, is composed of smaller EVs, which are less heterogenous in size, and that are marked by TET8^1^. We first characterized the APF fraction from IR-*CFA6*/*HRPL*#4 plants co-purified with PEN1-positive EVs. To do so, we vacuum-infiltrated IR-*CFA6*/*HRPL*#4 plants with Vesicle-Isolation Buffer (VIB, see methods), and further collected the apoplastic fluids (APFs) by low-speed centrifugation (at 900 g for 15 min, at 4°C). To remove dead cells and cell debris, these APFs were filtered (using a 0.2 µm filter), and the resulting filtered APFs further subjected to ultracentrifugation (at 40,000 g for 60 min, at 4 °C), using a swing-out rotor (TST41.14). Nanoparticle tracking analysis (NTA) from the recovered “P40” pellets revealed the presence of particles in a size range between 50 to 400 nm (Supplementary Fig. 4a). Transmission Electron Microscopy (TEM) further unveiled particles exhibiting a cup-shaped structure and/or a round vesicle-like morphology, as previously described^32^, and their diameters ranged between 16 to 409 nm with a mean of 90 nm +/-4 nm (Supplementary Fig. 4b, c). Furthermore, and as previously shown^31^, the EV markers PEN1 and PEN3 were readily detected in these P40 pellets (Supplementary Fig. 4d). Endogenous sRNAs sequenced from these APF fractions were mostly composed of tiny RNAs (tyRNAs; 10-17 nt), and were deprived of typical 21 and 24 nt peaks (Supplementary Fig. 4e)^33,34^. The sRNAs that mapped to the IR-*CFA6*/*HRPL* hairpin were composed of both 21 nt sRNAs and tyRNAs (Fig. 4a), and their accumulation profiles along the *CFA6*/*HRPL* regions of the hairpin were distinct from the ones of total sRNAs (Fig. 1e, 4b). The latter observation suggests that extracellular sRNAs are either selectively released in this apoplastic fraction, and/or stabilized by specific RNA-binding proteins, as previously proposed^5,34^. Furthermore, these anti-*cfa6*/*hrpL* sRNAs are unlikely derived from degraded cells and/or debris from IR-*CFA6*/*HRPL*#4 leaves, because we were unable to detect the U6 small nuclear RNA (snRNA) in the P40 fractions (Fig. 4c), which is readily detectable from total RNAs (Fig. 2a, e; Supplementary Fig. 1b, c, f; Supplementary Fig. 3).

**Figure 4.**
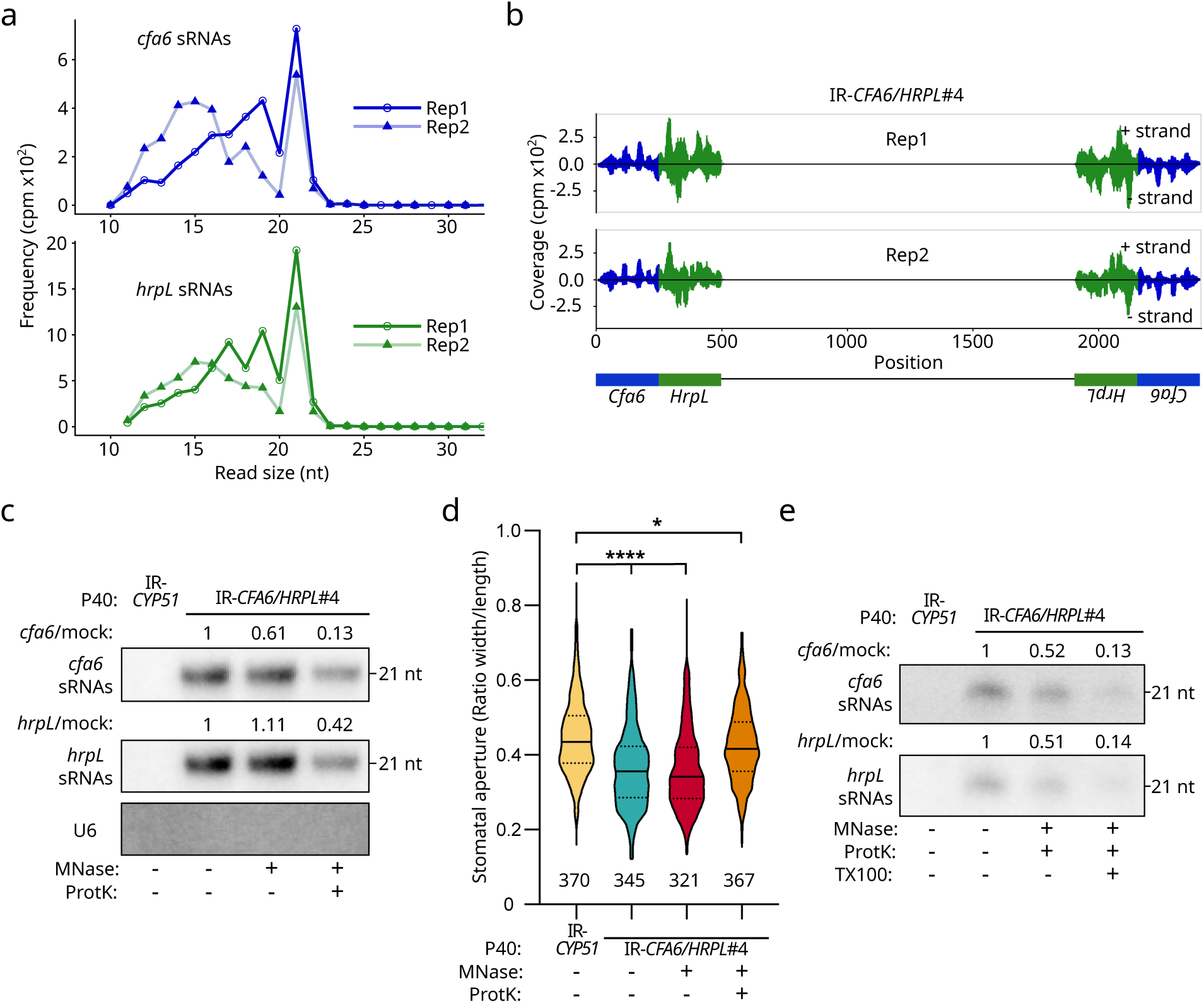
A pool of anti-*cfa6*/*hrpL* sRNAs, associated to proteins that are located outside EVs, suppresses *Pto* DC3000-triggered stomatal reopening. **a.** Size distribution and abundance of anti-*cfa6*/*hrpL* sRNA reads from MNase-treated P40 pellets of IR-*CFA6/HRPL*#4 plants. **b.** Small RNA reads from the P40 pellets of IR-*CFA6*/*HRPL*#4 plants that map to the *CFA6*/*HRPL* inverted repeat are depicted. Anti-*cfa6* and anti-*hrpL* sRNAs are shown in blue and green, respectively. **c.** Anti-*cfa6*/*hrpL* sRNAs accumulation is reduced when the P40 pellets of IR-*CFA6*/*HRPL*#4 plants were treated with MNase and ProtK. The P40 pellets from IR-*CFA6*/*HRPL*#4 plants were treated with mock, MNase or MNase plus ProtK, and the accumulation of anti-*cfa6*/*hrpL* sRNAs was analyzed by northern blot analysis. P40 pellets from untreated IR*-CYP51* plants were used as a negative control. The membrane was also probed with U6 to make sure that the P40 fractions were not contaminated by intracellular RNAs. The band intensities of *cfa6 /hrpL* siRNAs were quantified using ImageJ. Band intensities were normalized to the expression level of the mock condition. The results are shown above each blot. This is one representative experiment out of the three independent experiments performed. **d.** The ability of *Pto* DC3000 to reopen stomata is impaired in response to P40 pellets from IR-*CFA6*/*HRPL*#4 plants, and this effect is compromised in the presence of P40 pellets treated with MNase and Proteinase K (ProtK). The stomatal reopening assay was performed on Col-0 leaf sections incubated with IR-*CFA6/HRPL*#4 or IR*-CYP51-*derived P40 pellets that were subjected to different enzymatic treatments. Statistical significance was analyzed using a one-way ANOVA test (ns: p-value ≥ 0.05; *: 0.05 > p-value ≥ 0.01; ****: p-value < 0.0001). Similar results were obtained in three independent experiments and are pooled in a single plot. **e.** Anti-*cfa6* and anti-*hrpL* sRNAs accumulation is significantly reduced after MNase, ProtK and Triton X100 (TX100) treatments. Accumulation of sRNAs in IR-*CFA6/HRPL*#4- and IR-*CYP51*-derived P40 pellets was detected by northern blot analysis. The band intensities of *cfa6/hrpL* siRNAs were quantified using ImageJ. Band intensities were normalized to the expression level of the mock condition. The results are shown above each blot. This is one representative experiment out of the two independent experiments performed.

We next investigated whether the anti-*cfa6*/*hrpL* sRNAs from the P40 pellets could be functional. To this end, we made use of the previously described semi-*in vitro* stomatal reopening assay (Fig. 2), performed here with P40 pellets instead of RNAs. The P40 pellets from IR-*CFA6*/*HRPL*#4 APFs suppressed stomatal reopening triggered by *Pto* DC3000, compared to the P40 pellets from IR-*CYP51* APFs (Fig. 4d). This response was also detected upon incubation with the P40 pellets of IR-*HRPL*#4 APFs, but was almost fully abolished in the presence of *Pto*Δ*hrpL* mut *HrpL*, but not *Pto*Δ*hrpL* WT *HrpL* (Supplementary 4f). Collectively, these data not only provide evidence that anti-*cfa6*/*hrpL* sRNAs from the P40 pellets are biologically active, but also that the population of anti-*hrpL* sRNAs operates in a sequence-specific manner in *Pto* DC3000 cells.

The Arabidopsis apoplastic P40 pellets are composed of a mixture of particles, including PEN1-positive EVs^34^. To determine in which particles the active anti-*cfa6*/*hrpL* sRNAs could be present, we subjected the P40 pellets from IR-*CYP51* and IR-*CFA6*/*HRPL*#4 APFs to different enzymatic treatments and subsequently monitored their activities. It is noteworthy that none of these enzymatic treatments alters the morphology, the concentration nor the size distribution of apoplastic EVs (Supplementary Fig. 4a-c). The micrococcal nuclease (MNase) treatment, which degrades unprotected RNAs that are outside EVs and/or in contact with EVs, did not affect the ability of the P40 pellets to suppress stomatal reopening, nor the accumulation of anti-*cfa6*/*hrpL* sRNAs, as revealed by northern blot analysis (Fig. 4c, d). These data suggested that the active anti-*cfa6*/*hrpL* sRNAs from the P40 pellets are either inside EVs and/or outside EVs but associated with proteins, and thus protected from MNase-mediated degradation. Consistent with the latter hypothesis, we found that the combined MNase and Proteinase K treatment led to a decrease in the accumulation of anti-*cfa6*/*hrpL* sRNAs, and almost fully suppressed *Pto* DC3000-induced stomatal reopening (Fig. 4c, d; Supplementary Fig. 4g). An extra Triton X-100 (TX100) treatment, which disrupts EVs integrity^31^, led to an enhanced reduction in anti-*cfa6*/*hrpL* sRNA levels (Fig. 4e). This suggests the additional presence of intravesicular anti-*cfa6*/*hrpL* sRNAs that might mildly contribute to AGS. Altogether, these data indicate that active anti-*cfa6*/*hrpL* sRNAs from the P40 pellets are mainly associated with proteins that are located outside PEN1-positive EVs.

### The apoplasic fraction that co-purifies with TET8-positive EVs contains active anti-*cfa6*/*hrpL* sRNAs that are mainly located inside EVs

We next characterized the APF fraction from IR-*CFA6*/*HRPL*#4 plants co-purifying with TET8-positive EVs. To this end, we collected the supernatants from the above P40 pellets and subjected them to ultracentrifugation (at 100 000 g for 60 min, at 4°C), using the same swing-out rotor. The resulting “P100-P40” pellets were subsequently washed and analyzed microscopically, molecularly and functionally. An immunogold labeling approach, using native antibody recognizing the large extravesicular loop (the EC2 domain) of TET8 and a secondary antibody coating gold beads, confirmed the presence of TET8-positive EVs in these P100-P40 pellets (Fig. 5a). It is noteworthy that this subclass of EVs represented 22% of the whole vesicles analyzed from these APF fractions (Supplementary Fig. 5a). By contrast, none of the EVs from the P100-P40 fractions analyzed were labeled with an anti-UGPase antibody (Fig. 5a), indicating that the immunogold staining of TET8-positive EVs is specific. We next sequenced sRNAs from the P100-P40 pellets of IR-*CFA6*/*HRPL*#4 APFs. As observed with P40 pellets, endogenous sRNAs sequenced from P100-P40 pellets were mostly composed of tyRNAs (Supplementary Fig. 5b). Furthermore, the sRNAs that mapped to the *CFA6*/*HRPL* inverted repeat were composed of 21 nt sRNAs and tyRNAs (Fig. 5b), and their accumulation profiles along the *CFA6*/*HRPL* regions of the hairpin were distinct from the ones of total sRNAs (Fig. 1e, Fig. 5c). As previously observed with P40 fractions (Fig. 4c), we were unable to detect U6 snRNA from P100-P40 fractions (Fig. 5d), indicating that the detected anti-*cfa6*/*hrpL* sRNAs are unlikely derived from degraded cells and/or debris from IR-*CFA6*/*HRPL*#4 leaves.

**Figure 5.**
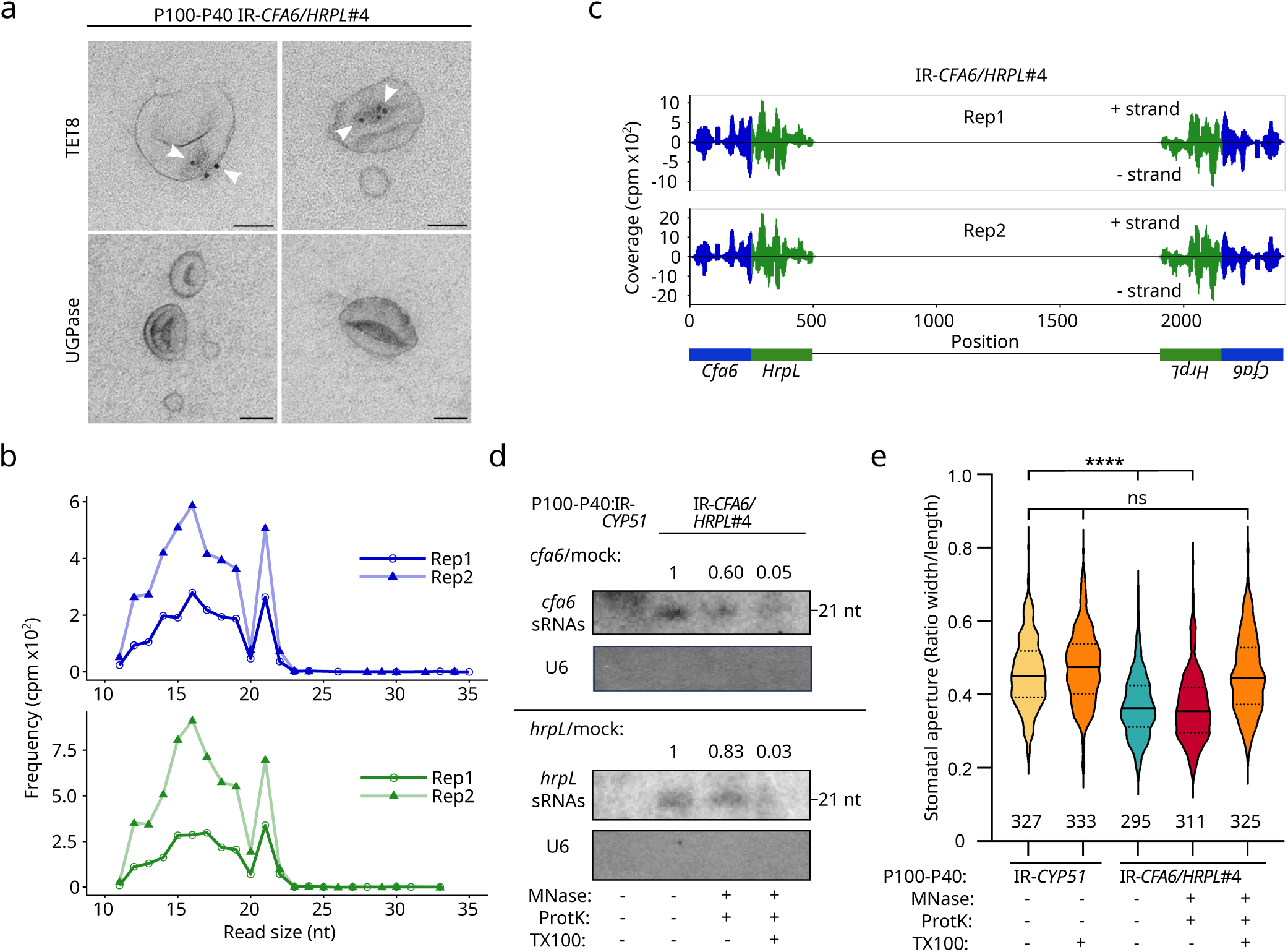
A pool of anti-*cfa6*/*hrpL* sRNAs, likely located inside TET8-positive EVs, suppresses *Pto* DC3000-triggered stomatal aperture. **a.** P100-P40 pellets contain TET8-positive EVs. EVs were visualized through TEM analysis, using native anti-TET8 (PhytoAB, PHY2750A) or UGPase (Agrisera) primary antibodies, followed by detection with a gold-conjugated secondary antibody. White arrows indicate the presence of gold beads. Scale bar: 50 nm. **b.** Size distribution and abundance of anti-*cfa6*/*hrpL* sRNA reads from MNase-treated P100-P40 pellets of IR-*CFA6/HRPL*#4 plants. **c.** Small RNA reads from the P100-P40 pellets of IR-*CFA6*/*HRPL*#4 plants that map to the *CFA6*/*HRPL* inverted repeat are depicted. Anti-*cfa6* and anti-*hrpL* sRNAs are shown in blue and green, respectively. **d.** Anti-*cfa6*/*hrpL* sRNAs accumulation is reduced when P100-P40 pellets of IR-*CFA6*/*HRPL*#4 plants were treated with TX100, MNase and ProtK. The P100-P40 pellets from IR-*CFA6*/*HRPL*#4 plants were treated with mock, MNase plus ProtK, or TX100 plus MNase plus ProtK and the accumulation of anti-*cfa6*/*hrpL* sRNAs was detected by northern blot analysis. P40 pellets from untreated IR*-CYP51* plants were used as a negative control. The membranes used were hybridized with either the *cfa6* probe (upper panel) or the *hrpL* probe (bottom panel). Of note, it was not possible to sequentially probe the same membrane with the *cfa6* and *hrpL* probes because of low abundance of these sRNAs from P100-P40 pellets. The band intensities of *cfa6/hrpL* sRNAs were quantified using ImageJ. Band intensities were normalized to the expression level of the mock condition. The results are shown above each blot. The membranes were also probed with U6 to make sure that the P100-P40 fractions were not contaminated by intracellular RNAs. **e.** The ability of *Pto* DC3000 to reopen stomata is impaired in response to P100-P40 pellets from IR-*CFA6*/*HRPL*#4 plants, and this effect is compromised when these P100-P40 pellets were treated with MNase, ProtK and TX100. Stomatal aperture measurements were performed on Col-0 leaf sections incubated with IR-*CFA6/HRPL*#4 or IR*-CYP51-*derived P100-P40 pellets that were subjected to different enzymatic treatments. Statistical significance was analyzed using a one-way ANOVA test (ns: p-value ≥ 0.05; ****: p-value < 0.0001). Similar results were obtained in three independent experiments and are pooled in a single plot.

We further subjected the P100-P40 pellets from IR-*CFA6*/*HRPL*#4 APFs to different enzymatic treatments and monitored their activities. Of note, none of these enzymatic treatments markedly alter the size distribution of EVs from these APF fractions, as revealed by TEM analysis (Supplementary Fig. 5c). Interestingly, in contrast with the results obtained with P40 pellets, the combined MNase and Proteinase K treatment did not alter the biological activity of P100-P40 pellets (Fig. 4d, 5e), nor the treatments with MNase or Proteinase K alone (Supplementary Fig. 5d, e). The former combined treatment did not markedly alter the accumulation of anti-*cfa6*/*hrpL* sRNAs either (Fig. 5d). These data indicate that anti-virulence sRNAs from the P100-P40 pellets are protected from enzymatic degradation. They also suggest that these active sRNAs are possibly protected inside the lumen of EVs from this APF fraction. Consistent with this hypothesis, we found that a triple treatment with MNase, Proteinase K and TX100, strongly altered the accumulation of anti-*cfa6*/*hrpL* sRNAs (Fig. 5d). In addition, it fully abolished the ability of the IR-*CFA6*/*HRPL*#4 P100-P40 pellets to suppress *Pto* DC3000-induced stomatal reopening (Fig. 5e). Altogether, these data demonstrate that the APF fraction co-purifying with TET8-positive EVs contains active antibacterial sRNAs that are likely located inside EVs.

### The apoplastic fraction exhibiting a substantial reduction in EVs is composed of active extracellular free sRNAs

We next investigated whether active anti-*cfa6*/*hrpL* sRNAs could additionally be present in the APF fraction that does not co-purify with EVs. For this purpose, we collected the supernatants from the above P100-P40 pellets. The recovered SN fractions exhibited a substantial reduction in canonical extracellular particles, as observed by NTA (Supplementary Fig. 6a), and were notably deprived of the PEN1 and PEN3 markers (Supplementary Fig. 4d). RNAs from the SN fraction of IR-*CFA6/HRPL*#4 APFs were further precipitated and subjected to sRNA-seq. Small RNAs that mapped to the Col-0 genome were almost exclusively composed of tyRNAs (Supplementary Fig. 6b). Anti-*cfa6*/*hrpL* sRNAs were represented by small amounts of 21 nt sRNAs and large amounts of tyRNAs (Fig. 6a). However, a more pronounced accumulation of 21 nt anti-*cfa6*/*hrpL* sRNAs, compared to cognate tyRNAs, was detected by northern blot analysis (Fig. 6b, Supplementary Fig. 6c). This suggests that biases occurring during sRNA library preparation (e.g. adapter ligation), coupled with yet-unknown post-transcriptional sRNA modifications, might hamper the retrieval of all 21 nt long anti-*cfa6*/*hrpL* sRNAs from these SN fractions. As observed for P40 and P100-P40 fractions, we also found that the accumulation profiles of SN-derived anti-*cfa6*/*hrpL* along the *CFA6*/*HRPL* regions of the hairpin were different from the ones of total sRNAs (Fig. 1e; Fig. 6b). Furthermore, we were unable to detect U6 snRNA from SN fractions (Fig. 6b), indicating that the detected anti-*cfa6*/*hrpL* sRNAs are unlikely derived from degraded cells and/or debris from IR-*CFA6*/*HRPL*#4 leaves.

**Figure 6.**
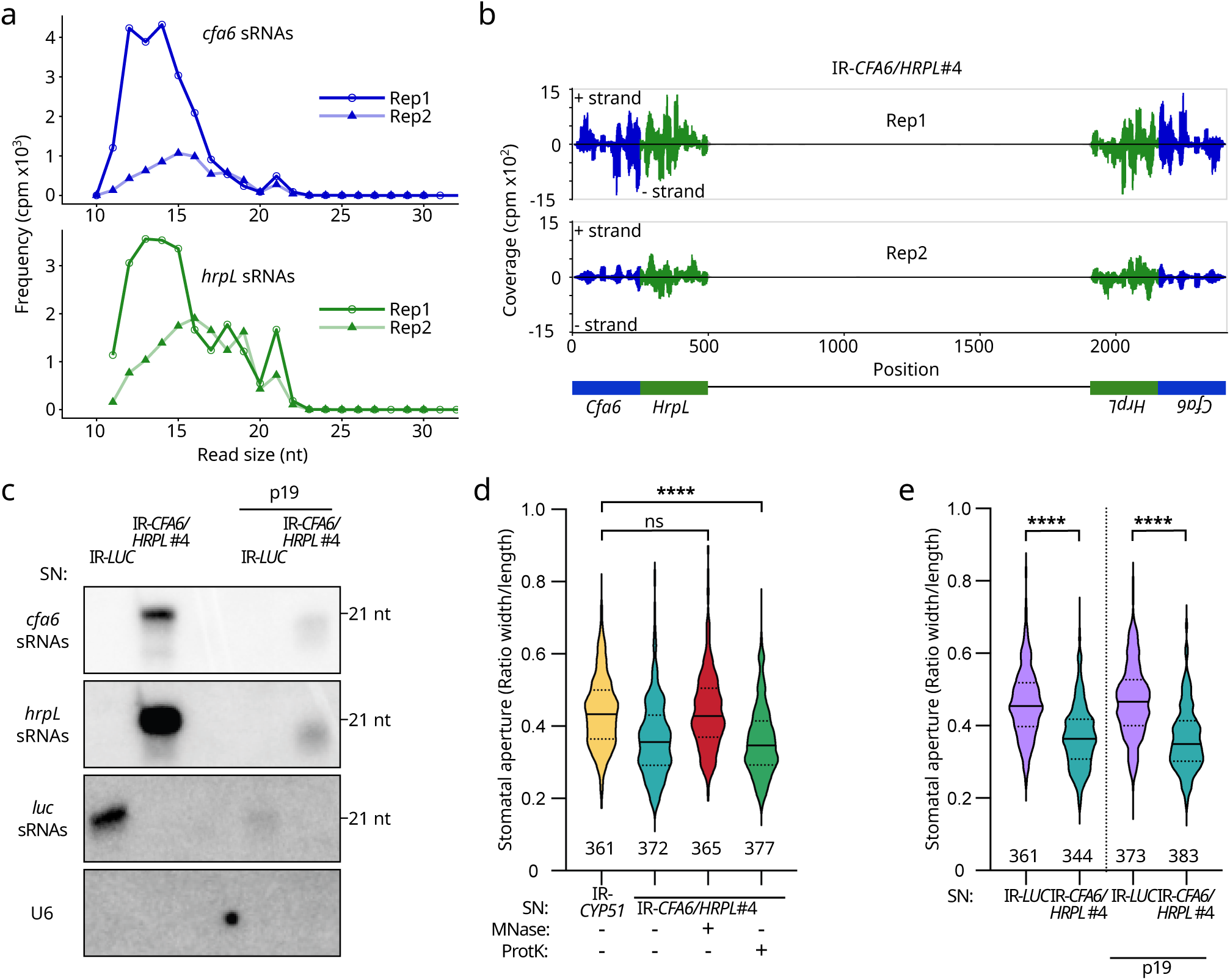
Extracellular free anti-*cfa6*/*hrpL* sRNA duplexes are recovered from SN fraction and harbour anti-virulence activity. **a.** Size distribution and abundance of anti-*cfa6*/*hrpL* sRNA reads from the SN fraction of IR-*CFA6/HRPL*#4 plants. **b.** Accumulation of anti-*cfa6*/*hrpL* sRNAs in IR-*CFA6/HRPL*#4 and IR-*LUC*-derived SN fractions before or after p19 pull-down assay was detected by northern blot analysis. U6 was used as a control of intracellular RNA. The membrane was also probed with U6 to make sure that the SN fractions were not contaminated with intracellular RNAs. This is one representative experiment out of the two independent experiments performed. **c.** Small RNA reads from the SN fraction of IR-*CFA6*/*HRPL*#4 plants that map to the *CFA6*/*HRPL* inverted repeat are depicted. Anti-*cfa6* and anti-*hrpL* sRNAs are shown in blue and green, respectively. **d.** The SN fraction from IR-*CFA6*/*HRPL*#4 plants prevents *Pto* DC3000-induced stomatal reopening, and this effect is abolished upon treatment of this APF fraction with MNase. Statistical significance was assessed using a one-way ANOVA test (ns: p-value ≥ 0.05; ****: p-value < 0.0001). Similar results were obtained in three independent experiments and are pooled in a single plot. **e.** P19 pulled-down sRNAs from the IR-*CFA6/HRPL*#4 SN fraction suppress *Pto* DC3000-induced stomatal reopening. The stomatal reopening assay was conducted as in c. Statistical significance was assessed using Student’s t-test (****: p-value<0.0001). Similar results were obtained in three independent experiments and are pooled in a single plot.

We further monitored the activity of the SN fraction from IR-*CFA6*/*HRPL*#4 APFs. This APF fraction was found to suppress stomatal aperture triggered by *Pto* DC3000, compared to the one derived from IR-*CYP51* plants (Fig. 6d). The same phenotype was detected in response to the SN fraction from IR-*HRPL*#4 APFs, but was impaired in the presence of *Pto*Δ*hrpL* mut *HrpL*, but not *Pto*Δ*hrpL* WT *HrpL* (Supplementary Fig. 6d). Collectively, these data indicate that anti-*cfa6*/*hrpL* sRNAs in the SN fraction are biologically active and that anti-*hrpL* sRNAs function in a sequence-specific manner in *Pto* DC3000 cells. To characterize in more depth the active sRNAs from the IR-*CFA6*/*HRPL*#4 SN fraction, we treated it with MNase or Proteinase K, and subsequently monitored its activity. The Proteinase K treatment did not alter the *Pto* DC3000-induced stomatal reopening phenotype (Fig. 6d), despite a substantial decrease in the accumulation of ∼21 nt anti-*cfa6*/*hrpL* sRNAs, and of the control microRNA miR167-5p (Supplementary Fig. 6c). Collectively, these results indicate that the majority of ∼21 nt anti-*cfa6*/*hrpL* sRNAs from the SN fraction is associated with proteins and not functional. To our surprise, we found that the MNase treatment alone fully abolished the ability of the IR-*CFA6*/*HRPL*#4 SN fraction to suppress *Pto* DC3000-induced stomatal reopening (Fig. 6d). Nevertheless, we were unable to detect, at the resolution of northern blot analysis, a significant decrease in the accumulation of anti-*cfa6*/*hrpL* sRNAs upon MNase treatment (Supplementary Fig. 6c). This effect was not due to a partial efficacy of the MNase treatment on sRNAs, because a full degradation of anti-*cfa6*/*hrpL* sRNAs was observed when RNA extracts from IR-*CFA6*/*HRPL*#4 plants were treated with this enzyme (Supplementary Fig. 6e). Collectively, these data suggest that a low-abundant pool of active anti-*cfa6*/*hrpL* sRNAs, and/or discrete sRNA species from the IR-*CFA6*/*HRPL* hairpin, must be unbound to proteins in the SN fraction, and thus sensitive to MNase action. We named these novel and functional extracellular sRNA species “extracellular free sRNAs” or efsRNAs.

### Anti-*cfa6*/*hrpL* efsRNA duplexes are recovered in the apoplast and are biologically active

The fact that i) anti-*cfa6*/*hrpL* duplexes, purified from IR-*CFA6*/*HRPL*#4 plants or *in vitro*-synthesized, were found functional (Fig. 2e-h), and ii) sRNA duplexes are typically more stable than single-stranded RNAs^35^, prompted us to investigate whether active anti-*cfa6*/*hrpL* efsRNA duplexes could be present in the apoplast. To purify sRNA duplexes from SN fractions, p19-chitin magnetic beads were incubated with the SN fractions from IR-*LUC* and IR-*CFA6/HRPL*#4 APFs, and the eluted bound sRNA duplexes were further analyzed by northern blot. We consistently retrieved comparable amounts of anti-*cfa6*/*hrpL* and anti-*luc* sRNAs in the p19 pulled-down RNAs from IR-*CFA6/HRPL*#4 and IR-*LUC* SN fractions, respectively (Fig. 6b). These data demonstrate the presence of both anti-*cfa6*/*hrpL* and anti-*luc* efsRNA duplexes in these APF fractions. Importantly, the p19 pulled-down sRNAs from the IR-*CFA6/HRPL*#4 SN fraction specifically suppressed the ability of *Pto* DC3000 to reopen stomata, to the same extent as total sRNAs from this apoplastic fraction (Fig. 6e). Collectively, these data provide evidence that the SN fraction of IR-*CFA6/HRPL*#4 APFs contains active efsRNAs, which are, at least in part, in a free and dsRNA form.

### A subset of SN-derived anti-*hrpL* sRNAs, including a functional 21 nt sRNA, is internalized by *Pto* DC3000 cells

Although anti-*hrpL* efsRNAs were found to be causal for AGS (Supplementary Fig. 6d), it was not experimentally shown whether they could be transferred in the cytoplasm of *Pto* DC3000 cells. To test this, we incubated *Pto* DC3000 cells with SN fractions from IR-*CFA6/HRPL*#4 APFs for 2 hours. Bacterial cells were further collected, and their outer membrane lysed using a buffer containing Ethylenediaminetetraacetic acid (EDTA) prior to washing and RNA isolation (Fig. 7a)^36^. This stringent condition ensured that sRNAs adhering onto the bacterial surface were removed from our samples. Consistent with an effective degradation of the bacterial outer membrane, the EDTA treatment substantially increased the uptake of propidium iodide in *Pto* DC3000 cells (Supplementary Fig. 7a). Nevertheless, it also altered *Pto* DC3000 survival (Supplementary Fig. 7b), which could account for an under-representation of sequenced sRNAs inside bacterial cells. We then prepared sRNA sequencing libraries corresponding to mock- and SN-treated conditions. To retrieve anti-*hrpL* sRNAs that were specifically derived from these SN fractions, rather than from bacterial *hrpL* transcripts (e.g. RNA degradation products), we established a first bioinformatic filter. Briefly, this filter only keeps reads that were not present in the control samples, which were untreated with SN fractions. In comparison to anti-*hrpL* sRNAs derived from the SN fractions of IR-*CFA6*/*HRPL*#4 APFs (Fig. 6a), an over-representation of tyRNAs, in a 10-12 nt size window, was recovered inside *Pto* DC3000 (Fig. 7b, c). This suggests that *Pto* DC3000 either preferentially takes-up tyRNAs and/or that longer sRNAs are internalized by *Pto* DC3000 and further processed inside this bacterium. Nevertheless, we still detected a small peak of 21 nt long sRNAs that was constantly observed across the biological replicates (Fig. 7c). We further isolated the most represented 21 nt anti-*hrpL* internalized sequence in both replicates, and chemically-synthesized the corresponding sRNA duplex labeled with the fluorescent dye ATT0 565, positioned on the 5’ end of the sRNA strand that is complementary to the *HrpL* mRNA sequence (Fig. 7d, Supplementary Fig. 7c). Furthermore, because active anti-*hrpL* sRNAs were found to be dependent on DCLs for their biogenesis (Fig. 2a, b), we designed this synthetic sRNA duplex with 2 nt-3’ overhangs, to mimic a DCL product (Fig. 7d, Supplementary Fig. 7c). Confocal imaging of *Pto* DC3000-GFP cells incubated with this ATTO 565-labeled anti-*hrpL* sRNA duplex, confirmed its internalization by, and/or association with, bacterial cells (Fig. 7e, white arrows). Importantly, the cognate unlabeled sRNA duplex (WT *hrpL*) exhibited full functionality in a stomatal reopening assay, which was not the case of its cognate mutated version (mut *hrpL*), which behaved as a non-functional anti-*cyp51* sRNA duplex control (Fig. 7f). Altogether, these data demonstrate that anti-*hrpL* sRNAs are transferred from the SN fractions of IR-*CFA6*/*HRPL*#4 APFs to the cytoplasm of *Pto* DC3000 cells. They also show that a chemically-synthesized 21 nt sRNA duplex, corresponding to a *bona fide* internalized anti-*hrpL* sRNA, is functional inside *Pto* DC3000.

**Figure 7.**
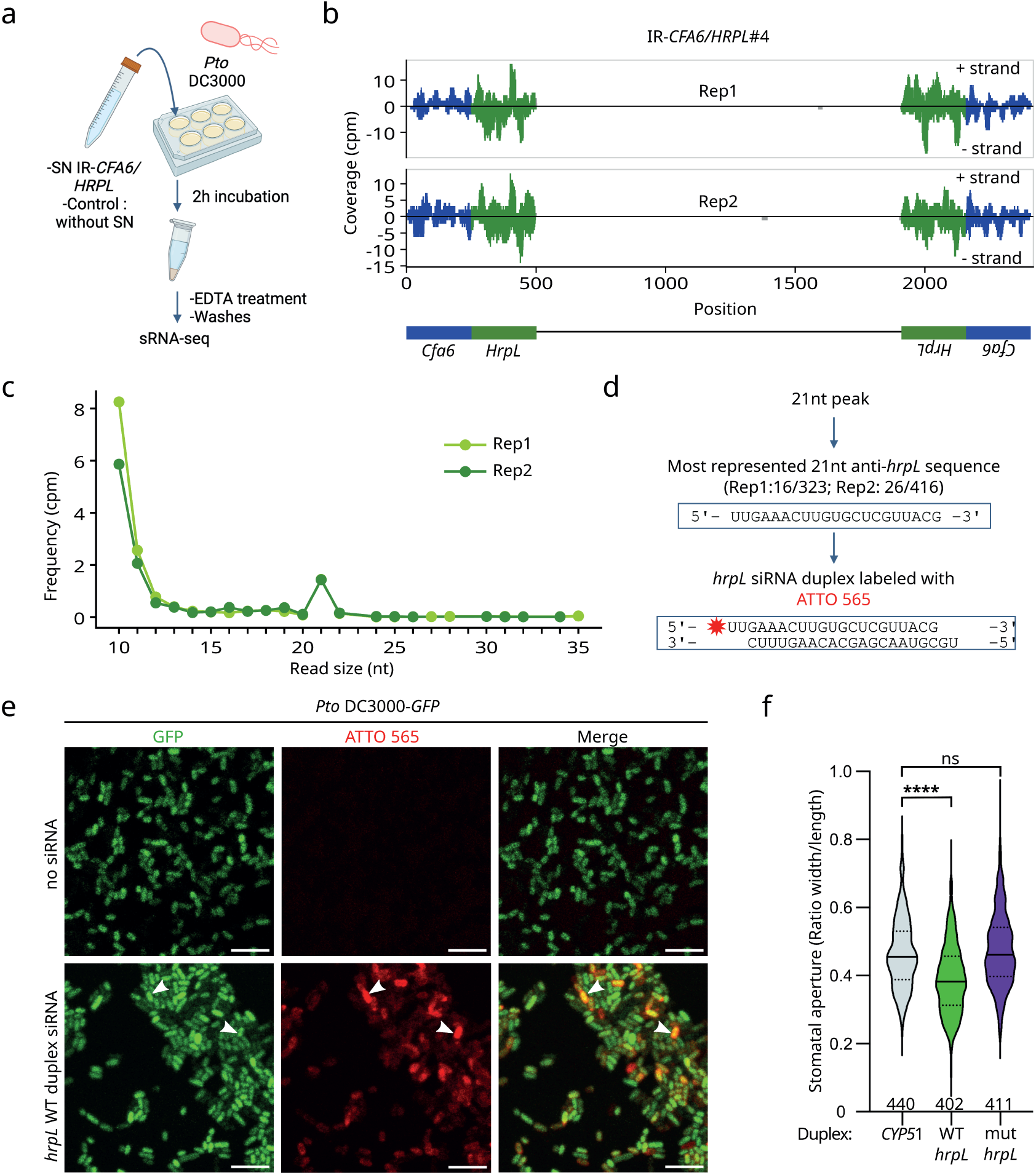
A subset of SN-derived anti-*hrpL* sRNAs is internalized by *Pto* DC3000 cells. **a.** Schematic representation of the experimental set-up employed to sequence sRNAs that are internalized by *Pto* DC3000. Bacterial cells were either incubated in VIB or in SN, derived from SA-treated IR-*CFA6/HRPL*#4 plants (see methods), for two hours before collection of the cells and washing with EDTA. Total RNAs were further extracted and used for sRNA-seq. **b.** Internalized sRNA reads that map to the *CFA6*/*HRPL* inverted repeat are depicted. Anti-*cfa6* and anti-*hrpL* sRNAs are shown in blue and green, respectively. **c.** Size distribution and abundance of anti-*cfa6*/*hrpL* sRNA reads. **d.** The most represented 21nt anti-*hrpL* sequence is framed in blue on top of the panel, with occurrence numbers for both replicates shown above it. Below is depicted the corresponding duplex version labeled with ATTO 565 (red star) in the 5’ end of the anti-*hrpL* sRNA strand that is complementary to *HrpL* mRNAs. **e.** Imaging of *Pto* DC3000-*GFP (*green) incubated with the *hrpL* duplex labeled with ATTO 565 (red). White arrows highlight examples of signals co-localization between ATTO 565 and GFP. Scale bar: 5 µm. **f.** WT *hrpL* duplexes but not *CYP51*, nor mut *hrpL* duplexes, suppress *Pto* DC3000-induced stomatal reopening. The stomatal aperture measurements were performed on Col-0 leaf sections incubated with sRNA duplexes. Statistical significance was analyzed using a one-way ANOVA test (ns: p-value ≥ 0.05; ****: p-value < 0.0001). Similar results were obtained in three independent experiments and are pooled in a single plot.

### Salicylic acid-treated IR-*CFA6*/*HRPL* plants trigger silencing of *HrpL* in *Pto* DC3000 cells

To monitor the silencing effect of anti-*hrpL* sRNAs on the *HrpL* target, we developed an experimental set-up, whereby *Pto* DC3000 cells expressing an epitope-tagged version of *hrpL* were exposed to extra-cellular/organismal sRNAs from IR-*CFA6*/*HRPL*#4 leaf sections (Fig. 8a). These conditions are reminiscent of the ones used in the stomatal reopening assay (Fig. 1f), with the exception that IR-*CFA6*/*HRPL*#4 plants were pre-treated with SA. It is noteworthy that SA promotes the expression and/or activity of RNAi factors^37–43^, as well as the release of Arabidopsis EVs, and of silencing factors, in the apoplast^31^, and was thus included here to eventually optimize the silencing effect mediated by anti-*hrpL* sRNAs. More specifically, IR-*CYP51* and IR-*CFA6/HRPL*#4 plants were pre-treated 48 hours before incubation with either mock or SA, and the corresponding leaf sections were further challenged for 2 hours with the *Pto* DC3000 strain expressing a functional C-terminal FLAG-tagged version of *hrpL* under its native promoter (Fig. 8a)^44^. The mRNA and protein levels produced by *HrpL* were subsequently monitored by RT-qPCR and western blot analyses, respectively. With this assay, we did not observe a consistent and significant decrease in the accumulation of *HrpL* mRNAs across different biological replicates, upon incubation of bacterial cells with mock or SA-pretreated IR-*CFA6*/*HRPL*#4 leaves (Supplementary Fig. 8). On the contrary, although leaf sections from IR-*CYP51* plants pretreated with SA exhibited a negative effect on HrpL protein accumulation, a more substantial reduction was always detected in the presence of IR-*CFA6/HRPL*#4 leaves treated in the same condition (Fig. 8b, c). Altogether, these data demonstrate that leaf sections of IR-*CFA6/HRPL*#4 plants pretreated with SA trigger the silencing of *HrpL* in *Pto* DC3000 cells. They also indicate that SA enhances HIGS efficacy from stable transgenic plants expressing antibacterial sRNAs.

**Figure 8.**
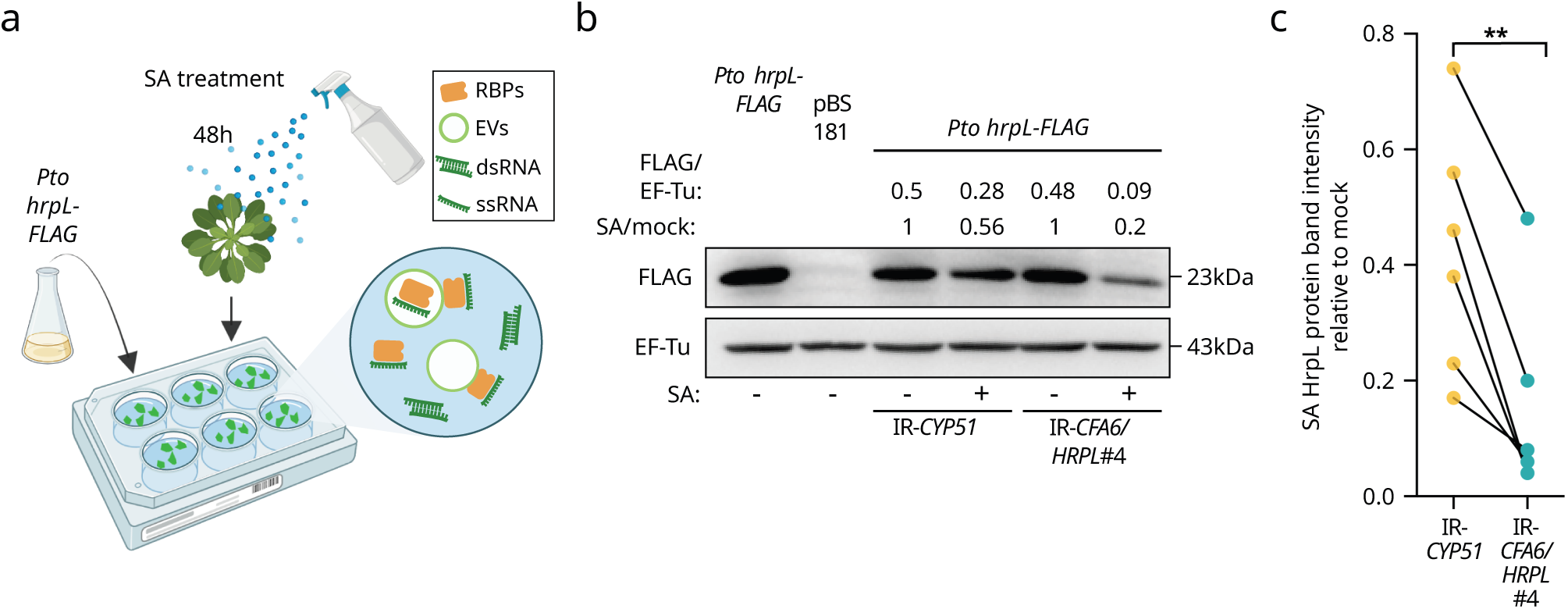
The accumulation of HrpL proteins is reduced in *Pto* DC3000 cells incubated with leaf sections of salicylic acid-treated IR-*CFA6*/*HRPL*#4 plants. **a.** Schematic representation of the experimental set-up employed to expose *Pto* DC3000 cells to plant-derived extra-cellular/organismal sRNAs. IR-*CFA6/HRPL*#4 or IR*-CYP51* plants were sprayed with SA (2 mM) for 48 hours prior to the experiment, and leaf sections from those plants were further incubated with a *Pto* DC3000 strain expressing a Flag-tagged version of *HrpL*, namely the *Pto* DC3000 *hrpL-FLAG* strain^44^. Some possible extra-cellular/organismal sRNA populations are illustrated in the circle. RBPs, EVs, dsRNA and ssRNA, stand for RNA-Binding Proteins, Extracellular Vesicles, double-stranded RNA and single-stranded RNA, respectively. **b**. Western blot analysis showing the reduction in the accumulation of HrpL proteins upon incubation of the *Pto* DC3000 *hrpL-FLAG* strain with leaf sections of SA-treated IR-*CFA6/HRPL*#4 plants, compared to leaf sections of IR-*CYP51* plants treated in the same condition. The accumulation of the elongation factor EF-Tu served as loading control. *Pto* DC3000 pBS181 contains the empty expression vector pBS181 and served as a negative control. This is one representative experiment out of the six independent experiments performed. **c.** Plot depicting the band intensity of *Pto* DC3000 *hrpL*-*FLAG* protein levels normalized to corresponding EF-Tu bands. IR-*CYP51* or IR-*CFA6/HRPL*#4 SA-treated ratios were then further normalized to IR*-CYP51* and IR-*CFA6*/*HRPL*#4 mock condition ratios, respectively. Similar results were obtained in six independent experiments and the data are pooled in a single plot. Statistical significance was assessed using Student’s paired t-test (ns: p-value ≥ 0.05).

### Endogenous sRNAs, from various genomic origins, are internalized by *Pto* DC3000 and exhibit predicted targets, some of which being repressed by SA biogenesis/signaling

We next assessed the potential contribution of endogenous Arabidopsis extracellular sRNAs on the targeting, and possibly silencing, of *Pto* DC3000 genes. To identify the putative *Pto* DC3000 targets of Arabidopsis sRNAs, we focused our analysis on SN-derived sRNAs that were found internalized by *Pto* DC3000 cells. For this purpose, we established a second bioinformatic filter, designed to retrieve, exclusively, Arabidopsis sRNAs inside *Pto* DC3000. Briefly, the sRNA sequences recovered from our first filter were selected, and those exhibiting multimapping between the Arabidopsis TAIR10 reference genome and the *Pto* DC3000 genome, further discarded. The size distribution analysis of these stringently filtered sRNAs unveils a majority of tyRNAs, which were derived from various genomic origins (Fig. 9a, b). This observation is consistent with an over-representation of this sRNA category from the SN-derived sRNAs mapping to the Arabidopsis genome (Supplementary Fig. 6b). It might additionally involve a further processing of SN-derived sRNAs inside bacterial cells. In agreement with the results obtained on transgenic lines expressing the artificial inverted repeat *CFA6*/*HRPL*, sRNAs produced from two well-described endogenous inverted repeats, namely IR71 and IR2039^45^, were also retrieved inside *Pto* DC3000 (Fig. 9b-f). However, in this case, only tyRNAs were recovered (Fig. 9c, d). The remaining internalized endogenous sRNAs were produced from other genomic regions, including mRNAs, transposable elements (TEs), rRNAs and tRNAs (Fig. 9b). In addition, because plant and mammalian tRNA-derived sRNAs (tsRNAs) i) are commonly retrieved in extracellular compartments and biofluids ^33,34,46,47^, and ii) have recently been shown to be transferred bidirectionally between host and microbes^17,48–52^, we investigated whether Arabidopsis tsRNAs could be part of the most represented sRNAs inside *Pto* DC3000. To do so, we extracted the top 100 most represented internalized sRNA sequences that are over 17 nt long in both replicates and examined the Arabidopsis tRNA database tRex^53^, for the presence of tsRNAs. We found that some of the top over-represented sRNA sequences are tRFs, derived from various tRNA precursors, including ProAGG.17 (Fig. 9g). Furthermore, we noticed that the corresponding tRF sequence displays plausible sequence-complementary targets in *Pto* DC3000 (Fig. 9h). Therefore, some Arabidopsis tRFs are taken-up by *Pto* DC3000 and exhibit a potential for targeting transcripts in this bacterium.

**Figure 9.**
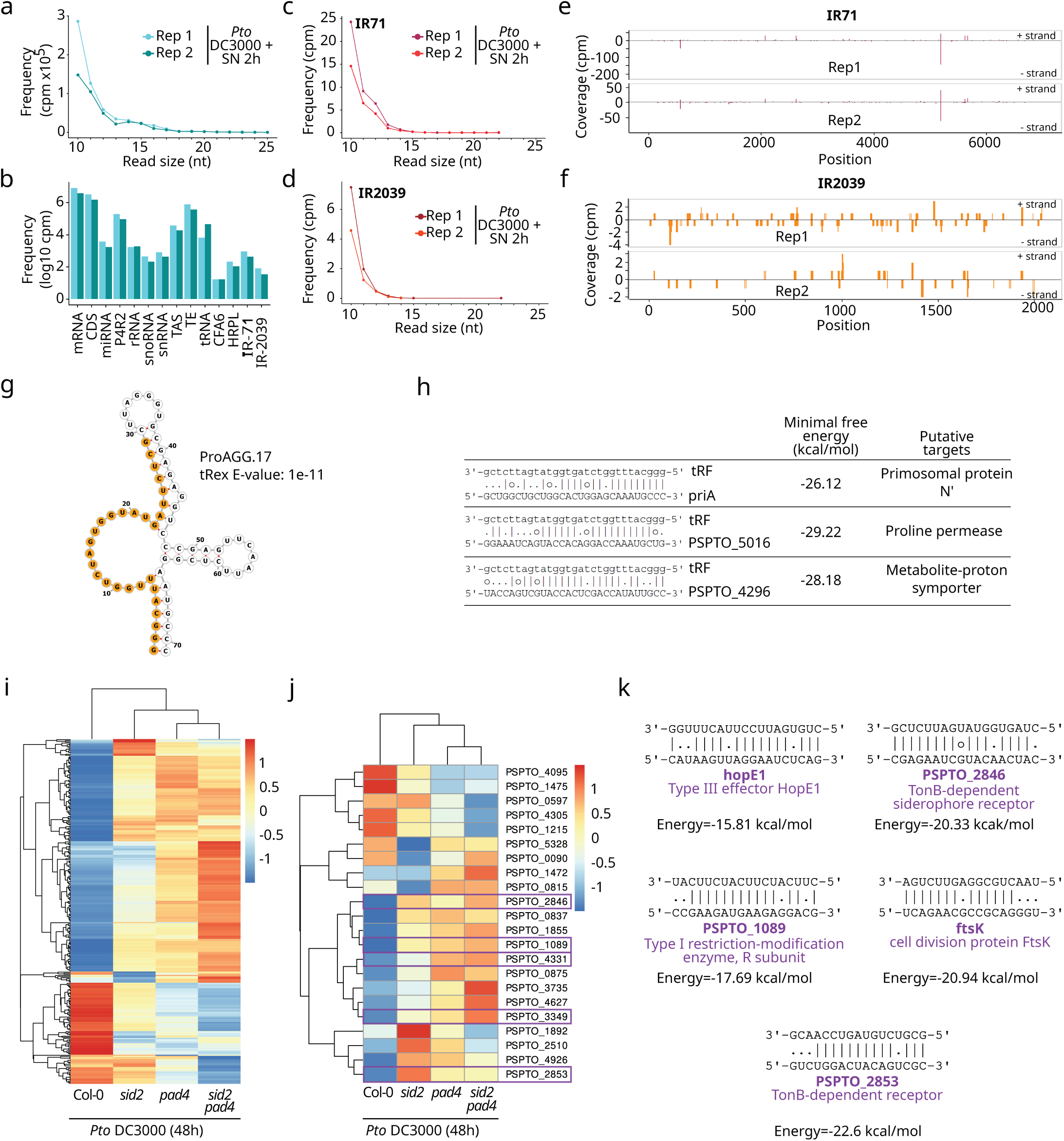
Endogenous sRNAs, from the SN of SA-treated IR-*CFA6/HRPL*#4 plants, are transferred into the cytoplasm of *Pto* DC3000 cells. **a.** Size distribution of all the endogenous sRNAs that were recovered in *Pto* DC3000 cells. SN fractions from SA-treated IR-*CFA6*/*HRPL*#4 plants were incubated for 2 hours with *Pto* DC3000 cells (*Pto* DC3000 + SN 2h), and the internalized endogenous sRNAs analyzed using the second stringent filter (see methods). **b.** Class repartition of reads after the second filtering step. **c.** Size distribution of reads mapping to IR71 after the second filtering step. **d.** Size distribution of reads mapping to IR2039 after the second filtering step. **e, f.** Coverage of sRNAs on IR71 (e) and IR2039 (f) after the second filtering step. **g.** RNA-fold structure of ProAGG.17. The most represented tRNA sequence from our filtered samples is highlighted in orange. **h.** Examples of predicted *Pto* DC3000 targets of the tRF highlighted in g. The three best predictions according to the minimal free energy value are represented. **i.** Three hundred seven *Pto* DC3000 SA-responsive proteins were predicted to be targets of internalized sRNAs. The heatmap depicts their accumulation in Col-0, *sid2*, *pad4* and *sid2 pad4* that were infected for 48 hours with *Pto* DC3000^54^. Z-score normalized (by row) values based on protein accumulation values reported in Nobori *et al*.^54^ **j.** Twenty-two of the SA-responsive proteins corresponded to targets of the top 100 most represented internalized sRNA reads. Their accumulation in Col-0-, *sid2-*, *pad4*- and *sid2 pad4*-infected plants is shown in the heatmap. **k.** Representation of some candidate sRNA-Target pairings corresponding to internalized sRNAs predicted to target mRNAs, whose protein levels are down-regulated in Col-0-infected plants but no longer repressed in *sid2*-, *pad4*-, and/or *pad4*/*sid2*-infected plants (from the 5 purple framed candidates depicted in j.).

Finally, because SA was found here to promote AGS (Fig. 8), we analyzed whether SA biogenesis and/or signaling contribute to the possible silencing of sRNA predicted targets in *Pto* DC3000. To do so, we made use of *Pto* DC3000 proteomic datasets generated from infected Col-0, the SA biogenesis mutant *sid2*, the SA signaling mutant *pad4*, and the double *sid2 pad4* mutant^54^. By cross-referencing the list of predicted *Pto* DC3000 targets from the whole set of internalized sRNAs, with the one of SA-responsive proteins, we found that 209 (68.1%), 213 (69.4%), 214 (69.7%), out of the 307 SA-responsive predicated sRNA targets, were less expressed in Col-infected plants compared to *sid2*-, *pad4*- and *sid2 pad4*-infected plants, respectively (Fig. 9i, Supplementary Table 4). A similar proportion of under-expressed targets, represented by 63.6% in Col-0 vs. *sid2*, 68.2% in Col-0 vs. *pad4* and 68.2% in Col-0 vs. *sid2 pad4*, was observed when the same analysis was performed on the predicted targets of the top 100 most represented internalized sRNAs, which are above 17 nt in size (Fig. 9j). Among those, we recovered functionally relevant *Pto* DC3000 genes, whose cognate mRNAs exhibit plausible Watson-Crick base-pairing with specific internalized sRNAs, including the type III effector HopE1, TonB dependent siderophore receptors, a type I restriction-modifying enzyme as well as the cell division-related gene *ftsk* (Fig. 9k). Altogether, these data indicate that the uptake of sRNAs by *Pto* DC3000 is not restricted to artificial sRNAs, but also occurs with endogenous sRNAs. They also highlight a potential for Arabidopsis endogenous sRNAs in targeting, and possibly silencing, *Pto* DC3000 genes, in part through SA biogenesis and/or signaling.

## Discussion

### Arabidopsis-encoded anti-*cfa6*/*hrpL* sRNAs are externalized from plant cells and dampen *Pto* DC3000 pathogenesis

We show here that Arabidopsis-encoded artificial sRNAs can target *Pto* DC3000 virulence-associated genes, resulting in the dampening of bacterial pathogenesis. In particular, we found that host-encoded sRNAs directed against the *Pto* DC3000 *Cfa6* and *HrpL* genes, fully suppressed bacterial-induced stomatal reopening (Fig. 1). Furthermore, we showed that, in Arabidopsis plants expressing anti-*hrpL* sRNAs, this phenotype no longer occurred in response to the *Pto*Δ*hrpL* mut *HrpL* strain (Fig. 3). Altogether, these findings indicate that antibacterial sRNAs act in a sequence-specific manner and can operate at the pre-invasive stage of infection, presumably by limiting COR biosynthesis in *Pto* DC3000 cells that come in contact with the leaf surface. In addition, we found that anti-*cfa6*/*hrpL* sRNAs reduced the ability of *Pto* DC3000 to multiply in the leaf apoplast of Arabidopsis upon dip-inoculation (Fig. 1). This phenotype might be caused by the inability of *Pto* DC3000 to reopen stomata in IR-*CFA6*/*HRPL* lines, thereby limiting its access to inner leaf tissues and thus to the apoplast, which is its main replicative niche. Collectively, these data imply that antibacterial sRNAs must be externalized from plant cells towards the surface of Arabidopsis leaves to reach epiphytic *Pto* DC3000 populations.

### Anti-*cfa6*/*hrpL* sRNA duplexes, but not their dsRNA precursors, are biologically active

By using genetic and molecular biology approaches (Fig. 2), we demonstrated that DCL-dependent antibacterial sRNAs, but not cognate dsRNA precursors, are the RNA entities responsible for AGS (Fig. 10). These data imply that *Pto* DC3000 must be capable of taking-up –passively and/or actively– anti-*cfa6*/*hrpL* sRNAs, despite the presence of a cell wall and an intricate double membrane structure. This assumption was confirmed by showing that artificial anti-*hrpL* sRNAs from the non-vesicular apoplastic fraction of IR-*CFA6*/*HRPL*#4 plants, but also a chemically synthesized 21 nt anti-*hrpL* sRNA duplex with 2-nt 3’ overhangs, were transferred into the cytoplasm of *Pto* DC3000 cells (Fig. 7). These results thus unveil major differences with environmental RNAi in eukaryotic organisms, such as *C. elegans* and plant herbivores, which specifically rely on long dsRNAs, or filamentous pathogens, which are dependent on both dsRNAs and sRNAs^55^. They also suggest that the industrial dsRNA production platforms, currently designed to produce long dsRNAs against pathogens and parasites, are not adapted to produce RNA-based biocontrol agents against bacterial pathogens.

**Figure 10.**
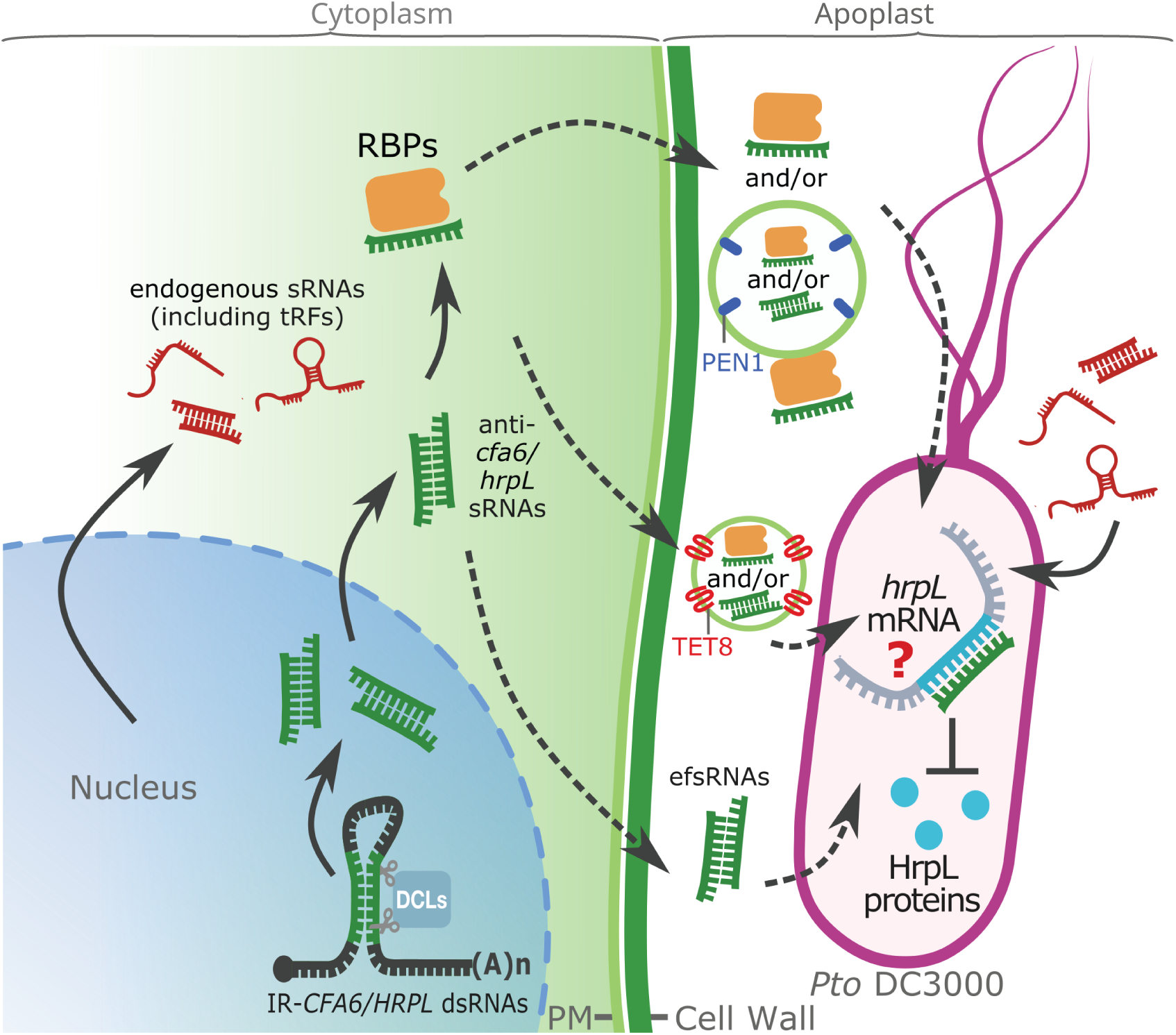
Model depicting the extracellular sRNA populations that are active against *Pto* DC3000 and/or internalized by these bacterial cells. Model depicting the characterized Arabidopsis-encoded artificial sRNA species directing gene silencing in interacting *Pto* DC3000 cells. The Arabidopsis IR-*CFA6/HRPL*#4 transgenic plants produce an inverted repeat transcript that is processed by DCL proteins into anti-*cfa6/hrpL* sRNAs (in green). The cytoplasmic anti-*cfa6/hrpL* sRNA species are further incorporated into protein complexes containing RNA Binding Proteins (RBPs), or potentially present in a free and dsRNA form in the cytosol, or in EVs. Both sRNA species are then exported in the apoplast through unknown mechanisms (dashed arrows). Three populations of active anti-*cfa6/hrpL* sRNAs have been characterized in this study: a first pool that is bound to proteins, which are either non-associated to PEN1-positive EVs, or potentially located on their surface. A second pool of sRNAs that is presumably located inside TET8-positive EVs, and a third pool that is unbound to proteins and in a free and dsRNA form, named efsRNAs. Active plant extracellular-sRNA species (*e*.*g*. anti-*hrpL* sRNAs), are subsequently transferred towards *Pto* DC3000 cells (dashed arrows), to direct sequence-specific gene silencing of *hrpL* through as-yet unknown mechanisms. Finally, the model depicts the endogenous sRNAs (in red), produced from various Arabidopsis endogenous genomic origins, including specific tRFs, which are internalized by *Pto* DC3000 cells.

### A pool of active apoplastic protein-bound sRNAs is located outside PEN1-positive EVs

We found here that a large proportion of apoplastic anti-*cfa6*/*hrpL* sRNAs is associated with proteins, and thus protected from MNase-mediated degradation (Fig. 4). This feature was notably observed for the apoplastic fraction that co-purified with PEN1-positive EVs (*i.e.* the P40 pellet), as well as the fraction exhibiting a substantial reduction in EVs and a depletion in PEN1 (*i.e.* the SN fraction). Nevertheless, only the protein-bound anti-*cfa6*/*hrpL* sRNAs from the P40 pellet, but not those from the SN fraction, were found active (Fig. 4; Fig. 6). In addition, protease and RNase protection assays from the P40 pellet revealed that these protein-bound antibacterial sRNAs are likely located outside PEN1-positive EVs. Based on these observations, we propose two possible non-mutually exclusive scenarios by which extravesicular protein-bound sRNAs from the P40 fraction could be transferred to, and/or effective in, *Pto* DC3000 cells (Fig. 10). The first one, which does not implicate EVs, would involve yet-unknown plant factor(s) from the protein-bound sRNAs complex(es) that would be specifically present in the P40 pellet, but not in the SN fraction. Potential candidates include the RNA-binding protein GRP7, as well as AGO2, which are both known to bind sRNAs^56–59^, and have been recovered in the apoplastic P40 fractions, in extravesicular forms^34^. The second one would involve PEN1-positive EVs, decorated with these protein-bound sRNAs complexes at their surface, which would act as carriers of these RNA-associated particles. The latter hypothesis is supported by the relatively large surface of EVs compared to their volume, which notably contains exposed phospholipids that can bind specific proteins from the extracellular environment^60^.

### A pool of active apoplastic sRNAs is likely located inside TET8-positive EVs

By characterizing the apoplastic fraction from IR-*CFA6*/*HRPL*#4 plants co-purifying with TET8-positive EVs (*i*.*e*. the P100-P40 fraction), we found that the MNase and Proteinase K combined treatment did not alter its antibacterial activity (Fig. 5). In contrast, an extra detergent treatment led to a substantial reduction in the accumulation of anti-*cfa6*/*hrpL* sRNAs, as well as a loss in AGS activity (Fig. 5). Collectively, these data indicate that a pool of active anti-*cfa6*/*hrpL* sRNAs is likely located inside TET8-positive EVs (Fig. 10). They are thus congruent with former studies showing that intravesicular plant antifungal sRNAs are delivered into *B*. *cinerea* through TET8-positive EVs^1,5^. They are also consistent with recent studies showing that i) antibacterial miRNAs, located inside ginger exosome-like nanoparticles (GELNs), can be delivered and active in the periodontal pathogen *Porphyromonas gingivalis* and the commensal intestinal bacterium *Lactobacillus rhamnosus*^61,62^, and (ii), let-7b-5p, embedded in EVs from human airway epithelial cells, is translocated in the opportunistic human pathogen *Pseudomonas aeruginosa*, to reduce its ability to form biofilm and to enhance its antibiotic sensitivity^36^. These findings also pave the way for the development of novel EV-based delivery systems for antibacterial sRNAs.

### A pool of active apoplastic sRNAs is unbound to protein and in a dsRNA form

Intriguingly, we found that anti-*cfa6*/*hrpL* sRNAs from the SN fraction harbour anti-virulence activity, which was found abolished upon MNase treatment alone (Fig. 6). Furthermore, by pulling-down anti-*cfa6*/*hrpL* sRNA duplexes from these apoplastic fractions, we showed that these extracellular sRNA species were functional in a stomatal aperture assay (Fig. 6). These data indicate that a third pool of anti-*cfa6*/*hrpL* efsRNAs, which is unbound to protein, and in a free and dsRNA form, is biologically active (Fig. 10). Although our findings do not exclude the possibility that extracellular single-stranded RNA species (ssRNAs) could also be present and functional in the apoplast, they suggest that the dsRNA structure of the above characterized efsRNAs might represent a structural feature that favors their stability in the apoplast. This hypothesis would provide an explanation for why artificial sRNA duplexes, exogenously applied on different plant species, were transmitted through, and stable in, xylem vessels, which are part of the apoplast^63^. It is also possible that yet-unknown post-transcriptional modifications of these functional efsRNAs contribute to their stability in the Arabidopsis apoplast.

### Salicylic acid-treated IR-*CFA6*/*HRPL* plants trigger *hrpL* silencing in *Pto* DC3000

To monitor the potential effect of anti-*hrpL* sRNAs on the silencing of the *HrpL* gene, we incubated the *Pto* DC3000 strain expressing an epitope-tagged version of *HrpL*, with leaf sections of IR-*CFA6*/*HRPL*#4 plants. Using this assay, we found a substantial reduction in the accumulation of HrpL proteins in *Pto* DC3000, which was specifically detected when the above transgenic plants were challenged with SA (Fig. 8). Our findings thus provide insights into strategies that can be further developed to improve HIGS efficacy. In addition, we observed that the levels of *HrpL* mRNAs were not consistently and significantly reduced in the presence of SA-pretreated IR-*CFA6*/*HRPL*#4 leaf sections across different biological replicates (Supplementary Fig. 8). Although these findings do not exclude the possibility that internalized anti-*hrpL* RNAs direct mRNA cleavage and/or degradation of *HrpL* mRNAs in *Pto* DC3000, they suggest that extra-cellular/organismal anti-*hrpL* sRNAs, produced from SA-challenged IR-*CFA6*/*HRPL* plants, predominantly dampen the translation of *HrpL* mRNAs in these experimental settings (Fig. 10). This regulatory process might involve the internalization by *Pto* DC3000 cells of vesicular and/or non-vesicular plant factors that are secreted in the apoplast in response to SA^34,64^, and/or SA-induced plant signals, which would enhance anti-*hrpL* sRNA-directed translational repression of *HrpL* mRNAs in *Pto* DC3000. Although the detailed mechanism remains unknown, various sRNA-directed translational inhibition processes are known to operate in animals, plants and bacteria^65–68,68–73^. For instance, some RNA-based antisense oligonucleotides (ASOs), which are in the same size range as tyRNAs, siRNAs and miRNAs, but also longer bacterial small non-coding RNAs, which are typically between 50 to 500 nt in size, have been shown to trigger translational inhibition of their mRNA targets through different mechanisms in prokaryotic cells ^68,73–75^. Altogether, these data suggest that *Pto* DC3000 must have evolved a machinery to take charge of the internalized plant sRNAs and direct gene silencing. It will thus be appealing to identify such machinery and elucidate the principles of plant sRNA target recognition and mode of action in bacterial cells.

### Endogenous efsRNAs, including tRNA-derived sRNAs, are taken-up by *Pto* DC3000 cells

By sequencing SN-derived sRNAs from IR-*CFA6*/*HRPL* plants that are transferred into the cytoplasm of *Pto* DC3000 cells, we did not only retrieve artificial sRNAs from the *CFA6*/*HRPL* hairpin but also Arabidopsis endogenous sRNAs. The latter sRNAs were derived from various genomic origins, including endogenous inverted repeats, mRNAs, TEs, rRNAs and tRNAs (Fig. 9; Fig. 10). Interestingly, among the most abundant sRNAs internalized by *Pto* DC3000 cells, we recovered tRFs, some of which exhibiting plausible bacterial mRNA targets (Fig. 9). Transfer RNA-derived sRNAs have previously been shown to be transferred bidirectionally in host-bacteria interactions^17,48–52^. More specifically, an abundant tRF was recovered inside outer membrane vesicles (OMVs) from *P*. *aeruginosa*, and transferred into human airway cells^48^. Furthermore, this tRF displays predicted targets encoding kinases from the lipopolysaccharide (LPS)-stimulated MAPK signaling pathway, and was shown to dampen the LPS-mediated induction of interleukin-8 (IL-8) secretion in primary human airway epithelial cells, as well as keratinocyte-derived chemokine production in mouse lung. Interestingly, tRFs produced from the rhizobium strain *B. japonicum* have also more recently been shown to be transferred inside soybean cells and further loaded into host AGO1, thereby silencing negative regulators of nodulation^17^. Inversely, two human 5’ tRNA halves from oral keratinocytes, were shown to be secreted outside these cells and further taken-up from the saliva by the Gram-negative pathobiont *Fusobacterium nucleatum*^49,50^. Importantly, these tsRNAs were able to reduce the growth of different *F*. *nucleatum* strains *in vitro*, presumably by interfering with the ribosome-related function of endogenous bacterial tRNAs^50^. Based on these collective findings, we speculate that the abundant Arabidopsis tRFs recovered inside *Pto* DC3000 might either silence sequence-complementary mRNA bacterial targets, and/or dampen protein biogenesis by acting as decoys of bacterial tRNAs. In addition, we found that a large set of Arabidopsis sRNAs recovered inside *Pto* DC3000 exhibits predicted bacterial mRNA targets, whose protein levels were reduced by SA signaling and/or biogenesis during infection (Fig. 9). Although these analyses have not been conducted in the natural host of *Pto* DC3000, they highlight a potential for endogenous plant sRNAs in directly reprogramming bacterial gene expression during infection. However, further studies are necessary to test whether plant sRNAs can indeed regulate gene expression in natural host-bacteria interactions, and also shape plant microbiomes composition as well as bacterial genomes evolution in these contexts.

## Supplementary figure legends

**Supplementary Figure 1. Characterization of IR-*CFA6*/*HRPL*, IR*-CYP51* and IR-*LUC* transgenic lines**

**a.** Representative pictures of five-week-old Arabidopsis Col-0 plants and of independent homozygous transgenic plants expressing the 35S::IR-*CYP51* or the 35S::IR-*CFA6/HRPL* transgenes. **b.** Accumulation of anti-*cfa6*/*hrpL* sRNAs from two-week-old Arabidopsis seedlings was detected by low molecular weight northern blot analysis. U6 was used as a loading control. **c.** Accumulation of anti-*cyp51* or anti-*luc* sRNAs respectively from the Arabidopsis plants IR-*CYP51*#1 or #2 and IR-*LUC* was detected by low molecular weight northern blot analysis. U6 was used as a loading control. IR-*CYP51* line #2 or IR-*LUC* were further used as reference lines in subsequent assays. **d.** Size distribution and abundance of sRNA reads from the IR-*CFA6/HRPL*#4 transgenic line. Data from two biological replicates are presented, and the same replicates are used in Fig. 1d. **e.** Thermodynamic energy analysis revealed that the free energy of sRNA-*Pto* DC3000 target interactions (Kcal/mol), which are almost exclusively composed of sRNA/*cfa6* and sRNA/*hrpL* pairs (see Table S2), is significantly lower than that from sRNA-Arabidopsis transcript interactions (see Table S1), indicating that off-targets in Arabidopsis are unlikely (see also Table S2 depicting the high e-values for all the interactions between the anti-*cfa6* and anti-*hrpL* sRNAs and Arabidopsis genes). **f.** Anti-*cfa6/hrpL* sRNAs accumulation remains unchanged upon bacterial dipping assay. Anti-*cfa6*/*hrpL* sRNAs were detected by low molecular weight northern blot analysis. U6 was used as a loading control. Anti-*cfa6/hrpL* sRNAs and U6 band intensities were measured with ImageJ. The ratio of sRNAs to U6 was calculated, and the results were normalized to the expression level of the mock condition. The results are shown above each blot.

**Supplementary Figure 2. Characterization of the RNA entities after size separation.**

**a.** Size distribution of total, long and sRNAs from IR-*CFA6/HRPL*#4 plants was obtained with an Agilent Bioanalyzer 2100 equipped with an RNA Nano chip. Low molecular weight RNA fractions are encircled for each sample. 18S and 25S ribosomal peaks are highlighted. **b.** Agarose gel picture of ethidium bromide-stained total, long and sRNAs from IR-*CFA6/HRPL#4* plants.

**Supplementary Figure 3. Molecular characterization of 35S::*HRPL* transgenic lines.**

Accumulation of anti-*hrpL* sRNAs was assessed by low molecular weight northern blot analysis using total RNA extracts from the Arabidopsis stable transgenic lines IR-*HRPL*#1 and #4, expressing the 35S::IR-*HRPL* transgene. U6 was used as a loading control.

**Supplementary Figure 4. Impact of MNase, Proteinase K, and MNase plus Proteinase K treatments on PEN1-positive EVs integrity, anti-*cfa6*/*hrpL* sRNAs accumulation and *Pto* DC3000-triggered stomatal reopening.**

**a.** Nanoparticle Tracking Analysis (NTA) showing that enzymatic treatments do not affect particles size nor abundance in P40 pellets. Size distribution and abundance of nanoparticles in IR-*CFA6/HRPL*#4-derived P40 pellets after different treatments; results from untreated (Mock), MNase-treated or MNase and Proteinase K (ProtK)-treated were similar. Similar results were obtained in three independent experiments. This is one representative experiment out of the three independent experiments performed. **b.** Transmission Electron Microscopy (TEM) analysis also revealed that enzymatic treatments do not alter neither the size nor the morphology of EVs recovered in P40 pellets. IR-*CFA6/HRPL*#4-derived P40 pellets were observed after different treatments: untreated, MNase-treated or MNase and ProtK-treated. Scale bar: 100 nm. **c.** Pictures of representative EVs in b. were measured using ImageJ software. The number of vesicles measured is written underneath each condition and statistical significance was assessed using the ANOVA test (ns: p-value ≥ 0.05). This is one representative experiment out of the two independent experiments performed. **d.** PEN1 and PEN3 protein accumulations were detected by western blot analysis in IR-*CFA6/HRPL*#4 total protein, P40 pellet or Supernatant fraction (SN). UGPase was used as a control in both western blot experiments. These are one representative experiment out of the three independent experiments performed. **e.** Size distribution and abundance of sRNA reads from IR-*CFA6/HRPL*#4 MNase-treated P40 pellets. Data from two biological replicates are presented, and the same replicates are used in Fig. 4b. **f.** The sRNAs from the P40 pellet of IR-*HRPL*#4 plants suppress *Pto* DC3000-induced stomatal aperture in a sequence-specific manner. Stomatal reopening measurements were conducted on Col-0 leaf sections incubated with IR-*HRPL*#4 or IR*-CYP51*-derived P40 pellets that were subjected to MNase treatment. The stomatal aperture assay was conducted as in Fig 2c. Similar results were obtained in three independent experiments and are pooled in a single plot **g.** Anti-*cfa6* and anti-*hrpL* sRNAs accumulation remains stable after MNase treatment but decreases after MNase plus ProtK treatment. Small RNAs band signal intensity was assessed using ImageJ. This molecular analysis was performed in three independent experiments. Signal was normalized relative to the untreated condition and statistical significance was assessed using Student’s *t*-test (ns: p-value ≥ 0.05, ****: p-value<0.0001).

**Supplementary Figure 5. Characterization of the P100-P40 pellet, anti-*cfa6*/*hrpL* sRNAs accumulation and impact of MNase and Proteinase K treatments on TET8-positive EVs integrity.**

**a.** Percentage of EVs in the P100-P40 sample labeled with TET8 antibody. Immunolabeling was performed using TET8 antibodies and secondary antibodies coupled with gold beads. EVs were observed through TEM. **b.** Size distribution and abundance of sRNA reads from IR-*CFA6/HRPL*#4 MNase-treated P100-P40 pellets. Data from two biological replicates are presented, and the same replicates are used in Fig. 5b. **c.** Pictures of representative EVs were measured using ImageJ software. The number of vesicles measured is written above each condition and statistical significance was assessed using the ANOVA test (ns: p-value ≥ 0.05; *: 0.05 > p-value ≥ 0.01). Similar results were obtained in three independent experiments and the data are pooled in a single plot. **d.** MNase treatment alone does not affect P100-P40 pellet activity against *Pto* DC3000. Stomatal aperture measurements were conducted in Col-0 pre-treated with IR-*CFA6/HRPL*#4*-* or IR-*CYP51*-derived P100-P40 pellets that were subjected to enzymatic treatments. Leaf sections were infected with the *Pto* WT strain. The number of stomata analyzed per condition is written underneath each condition and statistical significance was assessed using a one-way ANOVA test (****: p-value < 0.0001). Similar results were obtained in three independent experiments and the data are pooled in a single plot. **e.** ProtK treatment alone does not affect P100-P40 pellet activity against *Pto* DC3000. The stomatal reopening assay was performed as described in d. Similar results were obtained in two independent experiments and the data are pooled in a single plot.

**Supplementary Figure 6. Characterization of the apoplastic SN fractions of IR-*CFA6*/*HRPL*#4 and IR*-CYP51* plants.**

**a.** Size distribution and abundance of nanoparticles in IR-*CFA6/HRPL*#4 P40 pellets or SN fraction. The analyses were performed using Nanoparticle Tracking Analysis (NTA) method. Similar results were obtained in three independent experiments. This is one representative experiment out of the three independent experiments performed. **b.** Size distribution and abundance of sRNA reads from the SN fraction of the IR-*CFA6/HRPL*#4 transgenic line. Data from two biological replicates are presented, and the same replicates are used in Fig. 6c. **c.** Anti-*cfa6* and anti-*hrpL* sRNAs accumulation is reduced after ProtK but not after MNase treatments, at the resolution of northern analysis. Accumulation of sRNAs in IR-*CFA6/HRPL*#4- and IR*-CYP51-*derived SN fractions was detected by low molecular weight northern blot analysis. MicroRNA167 (miR167-5p) was used as an endogenous miRNA control. This is one experiment out of the two independent experiments performed. **d.** The sRNAs from the SN fraction of IR-*HRPL*#4 plants suppress *Pto* DC3000-triggered stomatal aperture in a sequence-specific manner. Stomatal aperture measurements were conducted on Col-0 leaf sections incubated with IR-*HRPL*#4 or IR-*CYP51*-derived SN fractions. Leaf sections were inoculated with indicated bacterial strains and the stomatal aperture measurements were performed as in Supplementary Fig. 3e. Similar results were obtained in three independent experiments and are pooled in a single plot. **e.** MNase treatments fully degrade anti-*cfa6* and anti-*hrpL* sRNAs from total RNA extracts from IR-*CFA6*/*HRPL*#4 plants. Ten micrograms of IR-*CFA6/HRPL*#4 total RNAs were treated with MNase and anti-*cfa6/hrpL* sRNAs were further detected by low molecular weight northern blot analysis. This is one representative experiment out of the three independent experiments performed.

**Supplementary Figure 7. Effect of EDTA treatment on *Pto* DC3000, and internalized anti-*cfa6/hrpL* 21nt long sRNAs.**

**a.** Treatment with EDTA effectively degrades the outer membrane of *Pto* DC3000. Membrane permeability was evaluated by measuring the internalization of propidium iodide (P. I) using fluorescence. The fluorescence signal was normalized to the optical density at OD at 600 nm (OD). **b.** *Pto* DC3000 exhibits impaired growth after treatment with EDTA. The bacteria were treated with EDTA to disrupt the outer membrane and periplasm, then cultured in NYGB medium for 24 hours. *In vitro* growth was monitored by measuring OD_600_. **c.** Upper panels: representations of the sRNA duplex with 2-nt 3’ overhangs for the WT *hrpL* duplex (*hrpL* WT sRNA duplex) depicted in green, and its cognate mutated version (*hrpL* mut sRNA duplex), depicted in purple. Anti-*hrpL* WT sRNA (shown in green) is 100% complementary to *HrpL* mRNA, whereas anti-*hrpL* mut sRNA (shown in purple) has been mutated to alter anti-*hrpL* sRNA pairing with the *HrpL* mRNA.

**Supplementary Figure 8. *HrpL* mRNAs are not consistently and significantly altered in *Pto* DC3000 cells in the presence of leaf sections from IR-*CFA6*/*HRPL*#4 and IR*-CYP51* plants pretreated with salicylic acid.**

RT-qPCR analyses depicting the levels of *HrpL* mRNAs in four independent experiments. The *HrpL* mRNA levels display slight or no significant changes in the presence of extra-cellular/organismal sRNAs from IR-*CFA6*/*HRPL*#4 leaf sections (with or without SA). Four independent biological replicates are represented. Levels of *HrpL* mRNAs in *Pto* DC3000 cells exposed to either IR*-CYP51* SA or IR-*CFA6*/*HRPL*#4 SA were normalized to IR*-CYP51* mock or IR-*CFA6*/*HRPL*#4 mock, respectively. Statistical significance was assessed using Student’s t-test (ns: p-value ≥ 0.05; *: 0.05 > p-value ≥ 0.01).

## Materials & Methods

### Plasmid construction

The IR-*CFA6/HRPL* construct is composed of 250 bp regions of *Pto* DC3000 genes, *cfa6* (1-250 nt) and *hrpL* (99-349 nt), aligned in sense and antisense directions, with the intron of the petunia chalcone synthase gene (*CHSA)* in between. The control vector construct IR-*CYP51* is designed to target conserved regions from *F. graminearum* that encompass *CYP51A*, *CYP51B* and *CYP51C* genes^6^, with the same *CHSA* intron. The IR-*CFA6/HRPL* and IR-*CYP51* constructs containing EcoRI and SalI sites at both extremities were synthesized by GenScript®, and inserted by restriction enzyme digestion into a modified pDONR221-P5-P2 vector carrying additional EcoRI and SalI sites to facilitate the insertion of these long-inverted repeats. The plasmids containing the 35S::IR-*CFA6*/*HRPL* and 35S::IR-*CYP51* constructs were obtained by a double recombination between pDONR221-P5-P2 carrying the inverted repeat sequences and pDONR221-P1-P5r carrying the 35S promoter sequence, in the pB7WG Gateway destination vector using LR Clonase Plus (Life Technologies). These plasmids were then introduced into the *Agrobacterium tumefaciens* C58C1 strain. The IR-*HRPL* construct, which is composed of the same 250 bp region of *hrpL* as in the IR-*CFA6/HRPL* construct, was recombined using GreenGate technology^76^ to generate the *35S_pro_*:IR-*HRPL* plasmid, which was then transformed in the Agrobacterium C58C1 strain. To generate the WT *hrpL* and the mut *hrpL* plasmids, the wild type (WT) *hrpL* sequence was amplified from the genomic DNA isolated from *Pto* DC3000, while the mutant *hrpL* sequence was amplified from a mutated sequence synthesized by GenScript®. These two sequences were further cloned into pDONR207 vector using BP Clonase (Life Technologies) and then introduced by recombination using LR Clonase (Life Technologies) into the pBS0046 destination vector, which carries a constitutive *NPTII* promoter. Specific primers used for the purpose of cloning are listed in Supplementary table 5. The Ubi.U4::IR-*HRPL* construct, was generated the same way than 35S::IR-*HRPL* but using Ubi.U4 promoter instead of 35S promoter. The second control vector IR-*LUC* construct was also generated using the GreenGate technology^76^, and is composed of a 250 bp region of the firefly luciferase under the control of 35S promoter.

### Plant material and growth conditions

Stable transgenic lines expressing 35S::IR-*CFA6/HRPL*, 35S::IR-*CYP51,* 35S::IR-*LUC* and 35S::IR-*HRPL*, and UbiU4::IR-*HRPL* constructs were generated by transforming Arabidopsis (Columbia-0 accession, Col-0) plants using the Agrobacterium-mediated-floral dip method^77^. Three independent Arabidopsis T4 transgenic lines expressing the IR-*CFA6/HRPL* hairpin, #*4*, #*5* and #*10*, two independent Arabidopsis T2 transgenic lines expressing the 35S::IR-*HRPL* hairpin, #*1* and #*4,* and one reference Arabidopsis T4 transgenic line expressing the IR-*CYP51* hairpin #*2*, one T3 IR-*LUC* line, one T3 Ubi.U4::IR-*HRPL* line and one Ubi.U4::IR-*HRPL dcl234* line, generated by crossing the Ubi.U4::IR-*HRPL* line with *dcl234* mutant were generated and used in our experiments. Sterilized seeds of Arabidopsis Col-0 and the selected homozygous transgenic lines were first grown for 12-14 days at 22°C/ 19°C (day/night) on plates containing ½ x MS medium (Duchefa), 1% sucrose and 0.8% agar (with or without antibiotic selection), or in soil, in an 8h photoperiod under a light intensity of 100 μE/m^2^/s. Seedlings were then pricked out to individual soil pots and grown in the same environmentally controlled conditions described above. Four-to six-week-old plants were used for all the experiments.

### Bacterial strains

The *Pto*Δ*cfa6*-GFP (*Pto* DC3118-GFP) strain is a gift from S. Y. He, while the *Pto*Δ*hrpL* strain is a gift from C. Ramos^78^. The *Pto hrpL-FLAG* and *Pto* pBS181 strains are gifts from M. J. Filiatrault and S. Cartinhour, respectively^44^. The *Pto*Δ*hrpL* strain expressing the *GFP* reporter gene was generated by transforming the bacterial mutant strain with the GFP-pPNpt Green plasmid by electroporation and then plating at 28°C on NYGA medium (5 g L^-1^ bactopeptone, 3 g L^-1^ yeast extract, 20 ml L^-1^ glycerol, 15 g L^-1^ Agar) containing gentamycin (1 µg mL^-1^) for selection. To generate the *Pto*Δ*hrpL* expressing WT *HrpL* or mut *HrpL* constructs, the *Pto*Δ*hrpL* strain was transformed by electroporation with the NPTII::WT-*HrpL* or NPTII::mut-*HrpL* plasmids. The bacteria were then plated at 28°C on NYGA medium with gentamycin (1 µg mL^-1^), to select the transformants that were confirmed by PCR using primers specific to *HrpL* gene.

### Stomatal aperture measurements

For each indicated genotype, 0.5 cm^2^ square sections were dissected from unpeeled leaves of 3 plants and immersed in mock solution (water) or in bacterial suspension at 10^8^ cfu mL^-1^. After 3 hours, unpeeled leaf sections were stained with 10 μg mL^-1^ propidium iodide (Sigma) and the abaxial surface was observed with a SP5 or SP8 laser scanning confocal microscope. The stomatal aperture (width relative to length ratio) was measured using ImageJ software for at least 90-110 stomata per condition. For RNA and vesicles treatments, the leaf sections were incubated for one hour with total RNAs (20 ng μl^-1^), P40 pellets, P100-P40 pellets and SN fractions extracted from specified genotypes, before incubation with the bacteria. In specified experiments, 1 μM of Coronatine (COR; Sigma) or *in vitro* synthesized sRNAs or long dsRNAs were supplemented to the bacterial suspension. Two to 5 independent experiments were pooled in each plot.

### Bacterial dipping assay

Three hours after the beginning of the night cycle in a growth chamber, three plants per condition were dip-inoculated with *Pto* DC3000 at 5 x 10^7^ cfu mL^-1^ supplemented with 0.02 % Silwet L-77 (Lehle seeds). Plants were then immediately placed in chambers with high humidity. At two day-post-inoculation, bacterial titers were measured for individual infected leaves (n=8), using a classical serial dilution method^79^.Three independent experiments were pooled in the same plot.

### Purification of Apoplastic Fluid (APF), Extracellular Vesicles (EVs) and Supernatant Fraction (SN)

APF extraction was done as previously described^31^. Briefly, 12 to 24 rosettes from 5 to 7-week-old IR-*CYP51*, IR-*LUC*, IR-*CFA6/HRPL*#4 or IR-*HRPL*#4 plants were vacuum-infiltrated with vesicle isolation buffer (VIB; 20 mM MES, 2 mM CaCl_2_, 0.1 M NaCl, pH 6.0). Rosettes were then placed inside a 20 mL needleless syringe that was inserted in a 50 mL Falcon tube and centrifuged at 900 g for 15 minutes at 4°C. The apoplastic fluid (APF) was collected and filtered through a 0.2 µm filter to discard cell debris. The APF was subjected to an ultracentrifugation step at 40,000 g for 60 min at 4°C giving rise to P40 pellets containing EVs. The supernatant from P40 pellets was subjected to another ultracentrifugation step at 100,000 g for 60 min at 4°C to collect the P100 pellet. At this step the supernatant contains the lowest amount of canonical EVs and is named the SN fraction. The P100 pellet, referred to as P100-P40, was resuspended in VIB for washing and then ultracentrifuged at 100,000g for 60 minutes at 4°C, and used for further analyses. The size and concentration of EVs from P40 or P100-P40 pellets and SN fractions were analyzed using the ZetaView NTA instrument (Particle Metrix). All ultracentrifugations were performed using a swing-out rotor (TST41.14).

### Treatments of P40, P100-P40 pellets and SN fractions

When indicated, the P40 pellets, the P100-P40 pellets or the SN fractions were treated with enzymes to digest either RNAs and/or proteins that were not protected by EVs. To eliminate RNAs, Micrococcal Nuclease (MNase) (Thermo Fisher Scientific) was used at 0.2 U μL^-1^ at 37°C for 15 min. The reaction was stopped with 20 mM EGTA. To digest proteins, Proteinase K (Thermo Fisher Scientific) was used at 2 U μL^-1^ at 37°C for 15 min. To digest both RNAs and proteins, MNase was first added to the sample for 15 min at 37°C, inactivated by addition of EGTA, and then Proteinase K was added for 15 min at 37°C. To disrupt EVs, they were incubated with 1% of Triton X-100 at room temperature for 1 hour, prior to enzymatic treatments. For all experiments, a negative control was performed by incubating samples at 37°C without enzyme during the same timeframe as for the enzymatic treatments.

### Extracellular Vesicles observation by Transmission Electron Microscopy (TEM)

Five μL of P40, P100-P40 pellets or SN fraction were deposited on copper formvar coated, carbonated grids (Electron Microscopy Sciences - EM-FCF200-CU-SB) and incubated for 20 min at room temperature. Grids were fixed afterwards in 2% paraformaldehyde/ 1% glutaraldehyde in 100 mM phosphate buffer pH 7.4 for 20 min at room temperature. The grids were washed 6 times in distilled water for 2 min. For embedding and contrast, grids were next incubated in a solution of 2% methylcellulose/ 1% uranyl acetate mixed at a 9:1 ratio for 7 min on ice in the dark. The excess of fluid from the grids was absorbed on a Whatman filter paper n°1. Grids were air-dried for 1 hour before storage. Observations were performed on a Transmission Electron Microscope (TEM) TECNAI 12BT 120 kV (ThermoFisher/FEI) equipped with a CCD Orius 1000 (Gatan) camera. Image acquisition was performed with DigitalMicrograph. EV diameters were measured using ImageJ software.

### Immunolabelling for Transmission Electron Microscopy (TEM)

Five μL of P100-P40 pellets were deposited on copper formvar coated, carbonated grids (Electron Microscopy Sciences - EM-FCF200-CU-SB) and incubated for 20 min at room temperature. Grids were fixed afterwards in 2% paraformaldehyde in 100 mM phosphate buffer pH 7.4 for 5 min at room temperature. The grids were washed 3 times in distilled water for 2 min, and next incubated in a solution of 50 mM ammonium chloride in 100 mM phosphate buffer pH 7.4 for 30 min at room temperature. The grids were washed 3 times in 0.1 % BSA-c (AURION) in 100 mM phosphate buffer pH 7.4 for 2 min each, and next incubated in the same buffer supplemented with primary antibody (either antibody against TET8 purchased from PhytoAB (PHY2750A), or anti-UGPase, purchased from Agrisera) at a 1:50 ratio for 60 min in a humid chamber at room temperature. The grids were washed 6 times in 0.1 % BSA-c in 100 mM phosphate buffer pH 7.4 for 2 min, then incubated in the same buffer with a 1:20 ratio of Goat anti-Rabbit (GAR) gold conjugate reagent (6 nm, AURION) for 60 min in a humid chamber at room temperature in the dark. The grids were washed 6 times in 0.1 % BSA-c in 100 mM phosphate buffer pH 7.4 for 2 min, then washed 3 times in 100 mM phosphate buffer pH 7.4 for 2 min. Grids were fixed afterwards in 2% glutaraldehyde in 100 mM phosphate buffer pH 7.4 for 5 min at room temperature. The grids were washed 6 times in distilled water for 2 min, and next incubated in a solution of 2% Methylcellulose/ 1% uranyl acetate mixed at a 9:1 ratio for 7 min on ice in the dark for embedding and contrast. The excess of fluid from the grids was absorbed on a Whatman filter paper °1. Grids were air dried for 3 hours before storage. Observations were performed on a Transmission Electron Microscope (TEM) TECNAI 12BT 120 kV (ThermoFisher/FEI) equipped with a CCD Orius 1000 (Gatan) camera. Image acquisition was performed with DigitalMicrograph.

### Separation of long and small RNA fractions

From 100 μg of total RNAs extracted with the Tri-reagent (Sigma, St. Louis, MO), long and small RNA fractions were separated using the mirVana miRNA isolation kit (Ambion®) according to the manufacturer’s instructions. The long and small RNA fractions were visualized by agarose gel electrophoresis and further analyzed using a Bioanalyzer 2100 (Agilent Technologies).

### *In vitro* synthesis of Inverted Repeat RNAs

*In vitro* synthesized RNAs were generated following the instructions of the MEGAscript® RNAi Kit (Life Technologies, Carlsbad, CA). Templates were amplified by PCR using gene specific primers containing the T7 promoter. PCR amplification was done in two steps with two different annealing temperatures to increase the specificity of primer annealing. After the amplification step, PCR products were purified by gel extraction using the PCR Clean-up kit (Macherey-Nagel) to eliminate any unspecific PCR products. To produce dsRNAs, the purified PCR products were used as templates for *in vitro* transcription performed according to the MEGAscript® RNAi Kit instructions. After purification with the filter cartridges, the corresponding dsRNAs were processed in 18-25 nt sRNA duplexes by incubation with ShortCut® RNase III (NEB, Ipswich, MA) for 20 min at 37°C. Small RNAs were then precipitated overnight at -20°C in presence of 3 volumes of 100% EtOH and 0.1 volume of 3 M NaCl, and then were washed once with 70% EtOH. Each step of this process was followed by gel electrophoresis (TAE 1X, 1% agarose gel for DNA amplification and 2% agarose gel for RNAs) to verify the quality of RNAs.

### RNA Gel Blot Analyses

Accumulation of low molecular weight RNAs was assessed by Northern blot analysis as previously described^80^. Total RNA was extracted using Tri-reagent (Sigma, St. Louis, MO) according to the manufacturer’s instructions and stabilized in 50% formamide. RNA gel blot analysis of low molecular weight RNAs was performed on 30 μg of total RNAs and as described previously^80^. Prior to RNA extraction from P40 or P100-P40 pellets, 600 to 700 μL of those fractions were separated in equal volumes, and each fraction was further treated (with mock or the enzyme of interest). The corresponding total RNAs were subsequently extracted as described for total RNA fraction. All the extracted RNAs were loaded on gel for further low molecular weight northern blot analyses as previously described^80^. Prior to RNA extraction from SN fraction, 12 mL of SN fraction were separated in 3 equal volumes, each fraction (4 mL) was further treated (with mock or the enzyme(s) of interest), and the corresponding RNAs were precipitated overnight at -20 °C using 400 μL NaCl 5 M, 12 mL EtOH 100%. RNAs from the resulting pellets were extracted as described before and loaded on gel for further low molecular weight northern blot analyses as previously described^80^. Two hundreds nanograms of total RNAs from IR-*CFA6*/*HRPL*#4 plants were also used as control in the northern blot experiments in parallel of the SN samples (as shown for instance in Fig. 4e). Detection of U6 RNA was used to confirm equal loading of intracellular RNAs. DNA oligonucleotides complementary to miR167-5p and U6 sequences were end-labeled with γ-^32^P-ATP using T4 Polynucleotide Kinase (PNK) (Promega) following the manufacturer’s instructions. To detect anti-*cfa6/hrpL* sRNAs, specific ^32^P-radiolabeled probes were generated by amplifying regions of 150 bp to 300 bp with gene specific primers (see Supplementary Table 5) and the amplicons were labeled by random priming (Prime-a-Gene® Labeling System, Promega). The indicated probes were hybridized to the same membrane by sequential rounds of probing and stripping.

### P19 pull-down assays to purify sRNAs duplexes

To pull-down sRNA duplexes, purified MBP-p19 proteins associated with chitin beads were used. For this purpose, 40 μg of total RNAs, or 2 mL of SN fraction, were resuspended in binding buffer at a final concentration of: 20 mM Tris-HCl pH 7.0, 100 mM NaCl, 1 mM DTT, 0.1 mg/mL BSA, 1X Protease Inhibitor Cocktail (Sigma Aldrich), and were then incubated with 100 units of purified MBP-p19-CBD proteins (gift from New England Biolabs) for 1 hour at room temperature. Forty μL of chitin magnetic beads (NEB) were first blocked overnight at 4°C in 200 μL of pre-treatment buffer (20 mM Tris-HCl pH 7.0, 100 mM NaCl, 1 mM EDTA, 1 mM DTT, 1 mg/mL BSA) before to be added to the sample and were then incubated with rotation for 1 hour at room temperature. To elute sRNA duplexes bound to p19, the beads were first washed 5 times in 500 μL of wash buffer (20 mM Tris-HCl pH 7.0, 100 mM NaCl, 1 mM DTT, 100 µg/mL BSA) and then resuspended in 30 μL of elution buffer (20 mM Tris-HCl pH 7.0, 100 mM NaCl, 0.5% SDS) and incubated for 10 min at 37°C, followed by 10 min at RT under stirring. The elution step was repeated twice and both elution fractions were pooled. Each pooled elution fraction was divided into two for the detection of anti-*cfa6*/*hrpL* sRNAs (Northern blot analysis) and for activity test (stomatal reopening assay).

### *In vitro* bacterial incubation with ATTO 565-labeled anti-*hrpL* sRNA duplexes

A *Pto* DC3000-GFP culture was grown to OD600nm=0.6. Bacterial cells were harvested by centrifugation (4000 rpm, 5 min, same settings for all centrifugations in this section), washed with Milli-Q water, and diluted to OD600nm=0.2. A 200 µl aliquot of this bacterial suspension was then incubated with 2 µM of anti-*hrpL*-ATTO 565 duplexes in a 24 well-plate at room temperature on an orbital shaker (heidolph polymax 1040). The incubation was carried out for 24 hours at 20 rpm. Following incubation, bacteria were collected by centrifugation and washed with Milli-Q water and resuspended in 500 µl of Milli-Q water. To cross-link the bacteria, 1% formaldehyde was added and incubated for 15 min at room temperature. The bacteria cells were again pelleted and washed with Milli-Q water and resuspended in 500 µL of Milli-Q water. A 1/10 dilution of the bacterial suspension was prepared for fluorescence analysis. GFP fluorescence was observed using a Leica SP8 laser scanning confocal microscope (excitation: 488 nm; emission: 500-550 nm), and ATTO 565 fluorescence was detected (excitation: 565 nm; emission: 580-630 nm).

### *In vitro* bacterial incubation with Salicylic-Acid treated Arabidopsis leaves

Five-week-old IR-*CYP51* and IR-*CFA6/HRPL*#4 transgenic plants were sprayed with either water (mock) or SA (2 mM) solution supplemented with 0.02% Silwett. For each condition, 48h after the spray, 8 pieces of 1 cm^2^ were cut from 2-3 leaves and incubated in 6-well plates with 2 mL of *Pto* DC3000 *hrpL-FLAG* at 5 x 10^7^ cfu mL^-1^ in *hrpL*-derepressing medium (HDM, 10 mM fructose, 50 mM potassium phosphate buffer, 7.6 mM (NH_4_)_2_SO_4_, 1.7 mM NaCl, pH 6.0)^81,82^ for 2 hours. Bacterial cells were then collected and *hrpL-FLAG* expression was assessed at the RNA level, by RT-qPCR analysis, and at the protein level, by western blot analysis.

### Incubation of bacteria with SN fraction and EDTA treatment

One mL of *Pto* DC3000 bacteria at OD600nm = 0.4 was incubated with 1 mL of SN from IR-*CFA6/HRPL#4* plants for 2 hours at room temperature. Then, bacteria were pelleted, washed with PBS, and resuspended in 400 µL of TSE buffer (0.2 M Tris-HCl pH 8.0, 0.5 M sucrose, 1 mM EDTA) + 100 µL of 0.5 M EDTA to lyse the outer membrane. After a 15 min incubation at room temperature, bacteria were pelleted and washed with PBS again before RNA extraction. To assess the effectiveness of EDTA treatment in disrupting the outer membrane, 20 µL of EDTA-treated bacterial suspension were diluted in a final volume of 200 µL H_2_O, containing 1 µL of propidium iodide. The fluorescence of the propidium iodide, bound to bacterial DNA, was measured using an excitation wavelength of 535 nm and an emission wavelength of 617 nm. To evaluate the effect of EDTA treatment on bacterial viability, the treated bacteria were resuspended in 500 µL of H_2_O. A 100 µL aliquot of the EDTA-treated bacteria was inoculated into 500 µL of NYGB medium in a 96-well plate, and optical density (OD600nm) was recorded 24 hours after inoculation.

### Bacterial RNA extraction and Real Time Quantitative PCR analysis

Total RNAs from bacterial cells were extracted with Tri-Reagent (Sigma, St. Louis, MO) according to the manufacturer’s instructions, followed by DNAse (RQ1, Promega) digestion at 37°C to remove the genomic DNA. One μg of total RNA was reverse-transcribed using RevertAid H minus Reverse Transcriptase (Thermo Fisher Scientific) and random hexamers according to the manufacturer’s instructions. Gene expression was quantified by Real Time-qPCR, using Takyon SYBR Green Supermix and gene specific primers with the following parameters: a 384-well optical reaction plate was heated at 95°C for 10 min, followed by 45 cycles of denaturation at 95°C for 10 s, annealing at 60°C for 20 s, and elongation at 72°C for 40 s. A melting curve was performed at the end of the amplification by steps of 1°C (from 95°C to 50°C). Expression was normalized to that of the bacterial gene *gyrA*. Primer sequences are listed in Supplementary table 5.

### Protein extraction and western blot analysis

Total protein extraction from bacterial cells or plant vesicles was done according to Hurckman & Tanaka^83^. Equal amounts of proteins were resolved on SDS-PAGE in Laemmli buffer and were then electroblotted on Immobilon®-P PVDF Membrane (Millipore). Membranes were blocked during 1 hour in a 5% milk solution in 1X PBS-Tween 0.1%, then incubated overnight at 4°C with specific antibodies. Antibodies against PEN3, UGPase and EF-Tu were purchased respectively from Agrisera and Hycult Biotech. Antibody against PEN1 was a gift from Prof. Thordal-Christensen. The next day, membranes were washed 3 times for 10 minutes in PBS-Tween 0.1% then incubated 2 hours at room temperature in PBS-Tween milk 5% with the secondary antibody. Membranes were washed again 3 times for 10 minutes in PBS-Tween 0.1% and revelation was performed with Chemicapt ImageQuant400 with Immobilon forte HRP substrate (Sigma).

### Bioinformatic analysis – Thermodynamic energy analysis

Unique reads mapping to *cfa6* and *hrpL* genes of *Pto* DC3000 from two sRNA libraries were used for all BLAST analyses (Supplementary Fig. 1e). The 500 most abundant anti-*cfa6* and anti-*hrpL* sRNA reads were selected and blasted against annotated Arabidopsis genes (TAIR10) and *Pto* DC3000 coding sequences (CDS) (-e-value 10, -word_size 4, -ungapped, - reward 1, -penalty -1)^84^. The free energy of binding for each read paired with its top target was calculated using RNAup^85^. In Fig. 3c unique reads mapping to the *HrpL* gene of *Pto* DC3000 in the two sRNA libraries presented in Fig. 1d were used for BLAST analysis against the sequences in Fig. 1c, and the thermodynamic energy analysis was performed as described earlier. Target prediction was made by aligning the reverse complement of each sRNA against the *Pto* DC3000 genome using blastn^86^ (version 2.12.0+, Parameters: -outfmt 6 -evalue 20 - - max_hsps 10000 -max_target_seqs 1000 -word_size 4 -ungapped -reward 1 -penalty -1). Alignments were processed to keep up to 5 mismatches, where wobble pairs (G-U) matches were considered as a partial (0.5) mismatch. Predicted energy of the RNA-RNA pair between the sRNA and the target sequence was calculated using RNAup ^85^.

### Small RNA sequencing

Small RNA libraries were constructed and sequenced from four- to five-week-old leaves of IR-*CFA6/HRPL*#4 plants. Raw reads have been deposited at the NCBI SRA under Bioproject (PRJNA587213). Libraries were mapped against the *Arabidopsis thaliana* genome (v TAIR10.1 GCF_000001735.4), the IR-*CFA6/HRPL* sequence, and the *Pto* DC3000 genome using ShortStack (v 3.8.4) and Bowtie2 (v 2.3.4) with parameters --mismatches 0 --dicermin 15 --dicermax 30 --bowtie_m 20 --mmap f^87^. Coverage of mapped loci was obtained with the Genomic Alignments package in R^88^. Unique reads mapping to *Cfa6* and *HrpL* in both replicates were extracted and quantified from ShortStack, and results were mapped using samtools^89^.

## Acknowledgements

We thank S.-H. He for providing the *Pto* DC3118-GFP strain, C. Ramos for providing the *Pto* DC3000 *hrpL* mutant strain, M. J. Filiatrault and S. Cartinhour for providing the *Pto* DC3000 *hrpL*-*FLAG* strain and *Pto* pBS181, H. Thordal-Christensen for providing PEN1 antibody, L. Martin-Jaular and F. Cocozza from C. Théry Lab for their help on the use of the NTA, H. Jin for providing an anti-TET8 antibody, J. Alfano for providing the anti-EF-Tu antibody, BioRender.com for some illustrations shown in the figures of this article, and members of the Navarro laboratory for critical reading of the manuscript.

## Author contributions

A.R., J.Z., M.S-R., O.T., L.N. designed research; A.R., J.Z., M.C., M.S-R., O.T., A.C. performed research; A.R., J.Z., M.C., M.S-R., O.T., A.E.F., V.M., L.L., generated resources; A.P-Q and L.C. conducted the bioinformatic analyses; A.R., J.Z., M.C., M.S-R., O.T., A.E.F., L.C., L.N. analyzed data; A.R. and L.N. wrote the manuscript; all the authors reviewed and edited the manuscript.

