## Supplementary Figures for "Vesicular and non-vesicular extracellular small RNAs direct gene silencing in a plant-interacting bacterium"

**Supplementary Figure 1**

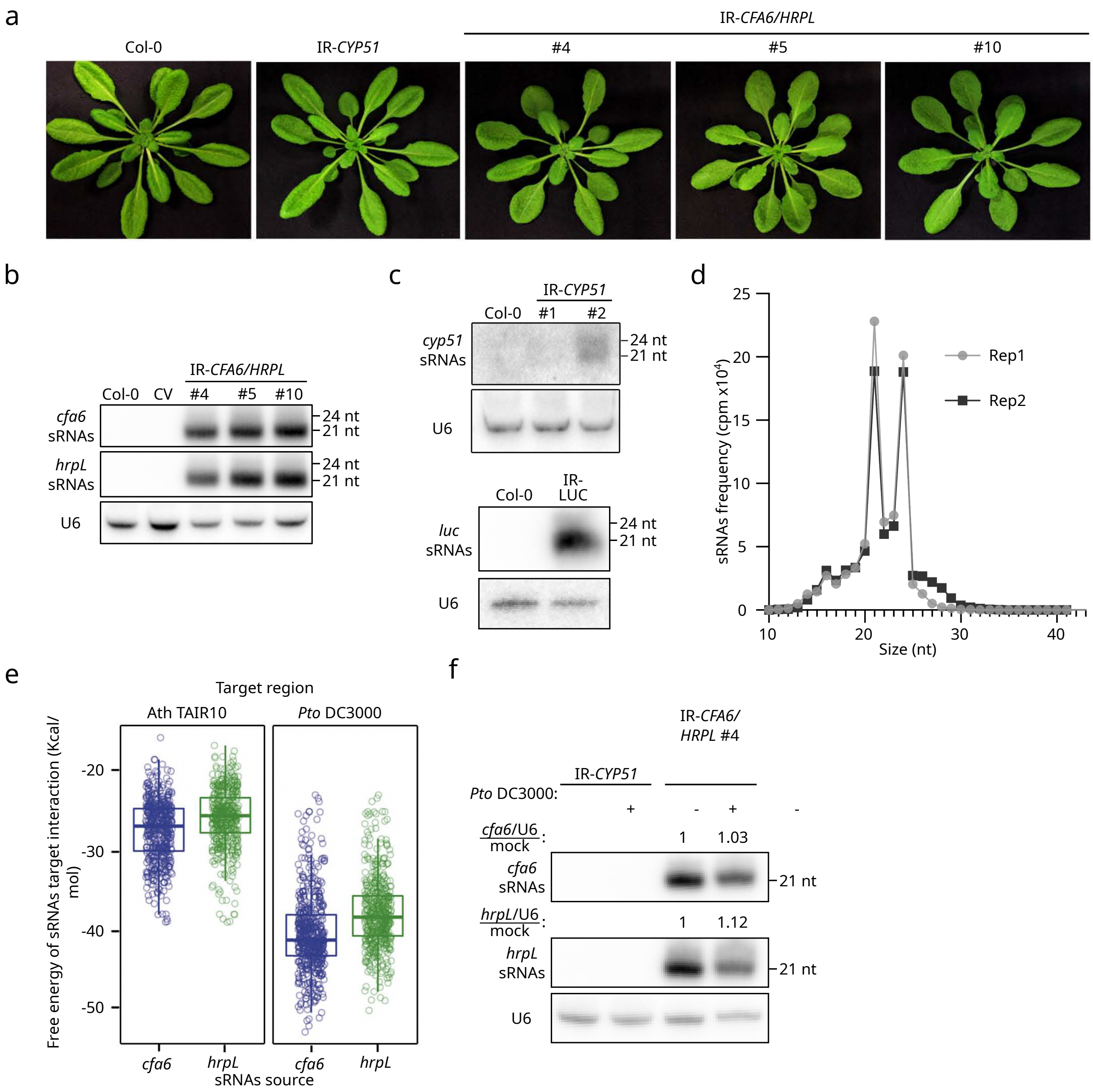

**Supplementary Figure 1. Characterization of IR-*CFA6/HRPL*, IR-*CYP51* and IR-*LUC* transgenic lines**

**a.** Representative pictures of five-week-old Arabidopsis Col-0 plants and of independent homozygous transgenic plants expressing the 35S::IR-*CYP51* or the 35S::IR-*CFA6/HRPL* transgenes. **b.** Accumulation of anti-*cfa6*/*hrpL* sRNAs from two-week-old Arabidopsis seedlings was detected by low molecular weight northern blot analysis. U6 was used as a loading control. **c.** Accumulation of anti-*cyp51* or anti-*luc* sRNAs respectively from the Arabidopsis plants IR-*CYP51*#1 or #2 and IR-*LUC* was detected by low molecular weight northern blot analysis. U6 was used as a loading control. IR-*CYP51* line #2 or IR-*LUC* were further used as reference lines in subsequent assays. **d.** Size distribution and abundance of sRNA reads from the IR-*CFA6/HRPL*#4 transgenic line. Data from two biological replicates are presented, and the same replicates are used in Fig. 1d. **e.** Thermodynamic energy analysis revealed that the free energy of sRNA-*Pto* DC3000 target interactions (Kcal/mol), which are almost exclusively composed of sRNA/*cfa6* and sRNA/*hrpL* pairs (see Table S2), is significantly lower than that from sRNA-Arabidopsis transcript interactions (see Table S1), indicating that off-targets in Arabidopsis are unlikely (see also Table S2 depicting the high e-values for all the interactions between the anti-*cfa6* and anti-*hrpL* sRNAs and Arabidopsis genes). **f.** Anti-*cfa6/hrpL* sRNAs accumulation remains unchanged upon bacterial dipping assay. Anti-*cfa6*/*hrpL* sRNAs were detected by low molecular weight northern blot analysis. U6 was used as a loading control. Anti-*cfa6/hrpL* sRNAs and U6 band intensities were measured with ImageJ. The ratio of sRNAs to U6 was calculated, and the results were normalized to the expression level of the mock condition. The results are shown above each blot.

**Supplementary Figure 2**

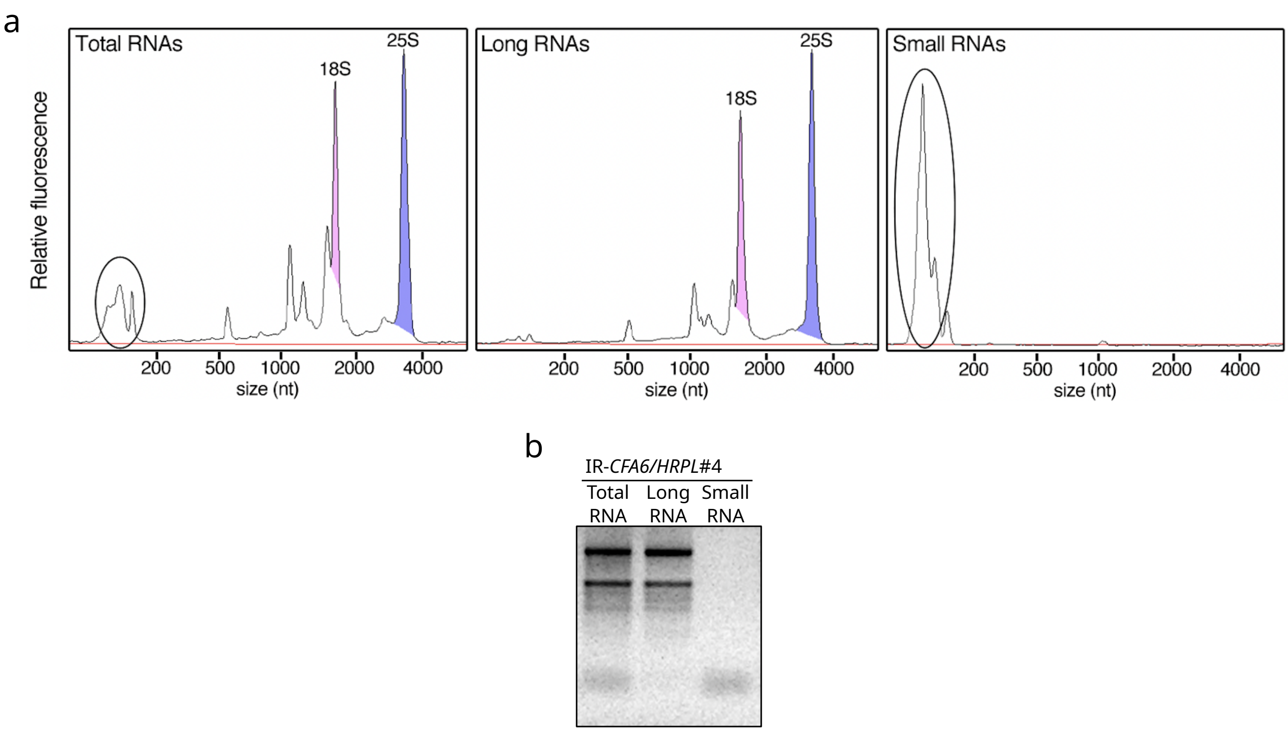

**Supplementary Figure 2. Characterization of the RNA entities after size separation.**
**a.** Size distribution of total, long and sRNAs from IR-*CFA6/HRPL*#4 plants was obtained with an Agilent Bioanalyzer 2100 equipped with an RNA Nano chip. Low molecular weight RNA fractions are encircled for each sample. 18S and 25S ribosomal peaks are highlighted. **b.** Agarose gel picture of ethidium bromide-stained total, long and sRNAs from IR-*CFA6/HRPL#4* plants.

**Supplementary Figure 3**

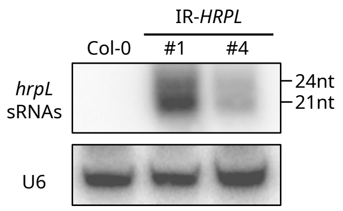

**Supplementary Figure 3. Molecular characterization of 35S::*HRPL* transgenic lines.** Accumulation of anti-*hrpL* sRNAs was assessed by low molecular weight northern blot analysis using total RNA extracts from the Arabidopsis stable transgenic lines IR-*HRPL*#1 and #4, expressing the 35S::IR-*HRPL* transgene. U6 was used as a loading control.

**Supplementary Figure 4**

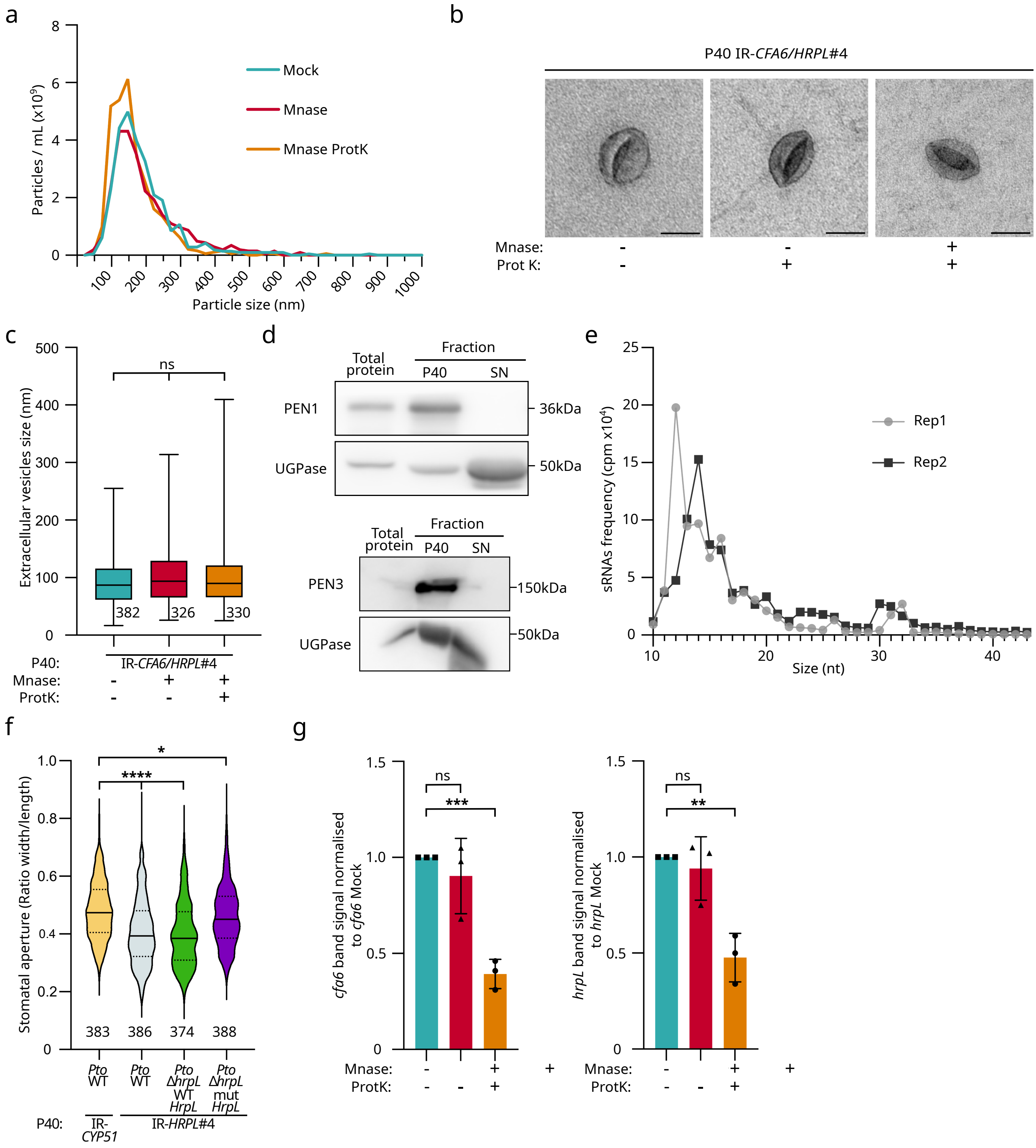

**Supplementary Figure 4. Impact of MNase, Proteinase K, and MNase plus Proteinase K treatments on PEN1-positive EVs integrity, anti-*cfa6*/*hrpL* sRNAs accumulation and *Pto* DC3000-triggered stomatal reopening.**

**a.** Nanoparticle Tracking Analysis (NTA) showing that enzymatic treatments do not affect particles size nor abundance in P40 pellets. Size distribution and abundance of nanoparticles in IR-*CFA6/HRPL*#4-derived P40 pellets after different treatments; results from untreated (Mock), MNase-treated or MNase and Proteinase K (ProtK)-treated were similar. Similar results were obtained in three independent experiments. This is one representative experiment out of the three independent experiments performed. **b.** Transmission Electron Microscopy (TEM) analysis also revealed that enzymatic treatments do not alter neither the size nor the morphology of EVs recovered in P40 pellets. IR-*CFA6/HRPL*#4-derived P40 pellets were observed after different treatments: untreated, MNase-treated or MNase and ProtK-treated. Scale bar: 100 nm. **c.** Pictures of representative EVs in b. were measured using ImageJ software. The number of vesicles measured is written underneath each condition and statistical significance was assessed using the ANOVA test (ns: p-value $\geq 0.05$). This is one representative experiment out of the two independent experiments performed. **d.** PEN1 and PEN3 protein accumulations were detected by western blot analysis in IR-*CFA6/HRPL*#4 total protein, P40 pellet or Supernatant fraction (SN). UGPase was used as a control in both western blot experiments. These are one representative experiment out of the three independent experiments performed. **e.** Size distribution and abundance of sRNA reads from IR-*CFA6/HRPL*#4 MNase-treated P40 pellets. Data from two biological replicates are presented, and the same replicates are used in Fig. 4b. **f.** The sRNAs from the P40 pellet of IR-*HRPL*#4 plants suppress *Pto* DC3000-induced stomatal aperture in a sequence-specific manner. Stomatal reopening measurements were conducted on Col-0 leaf sections incubated with IR-*HRPL*#4 or IR-*CYP51*-derived P40 pellets that were subjected to MNase treatment. The stomatal aperture assay was conducted as in Fig 2c. Similar results were obtained in three independent experiments and are pooled in a single plot **g.** Anti-*cfa6* and anti-*hrpL* sRNAs accumulation remains stable after MNase treatment but decreases after MNase plus ProtK treatment. Small RNAs band signal intensity was assessed using ImageJ. This molecular analysis was performed in three independent experiments. Signal was normalized relative to the untreated condition and statistical significance was assessed using Student's *t*-test (ns: p-value $\geq 0.05$, ****: p-value$<0.0001$).

### Supplementary Figure 5

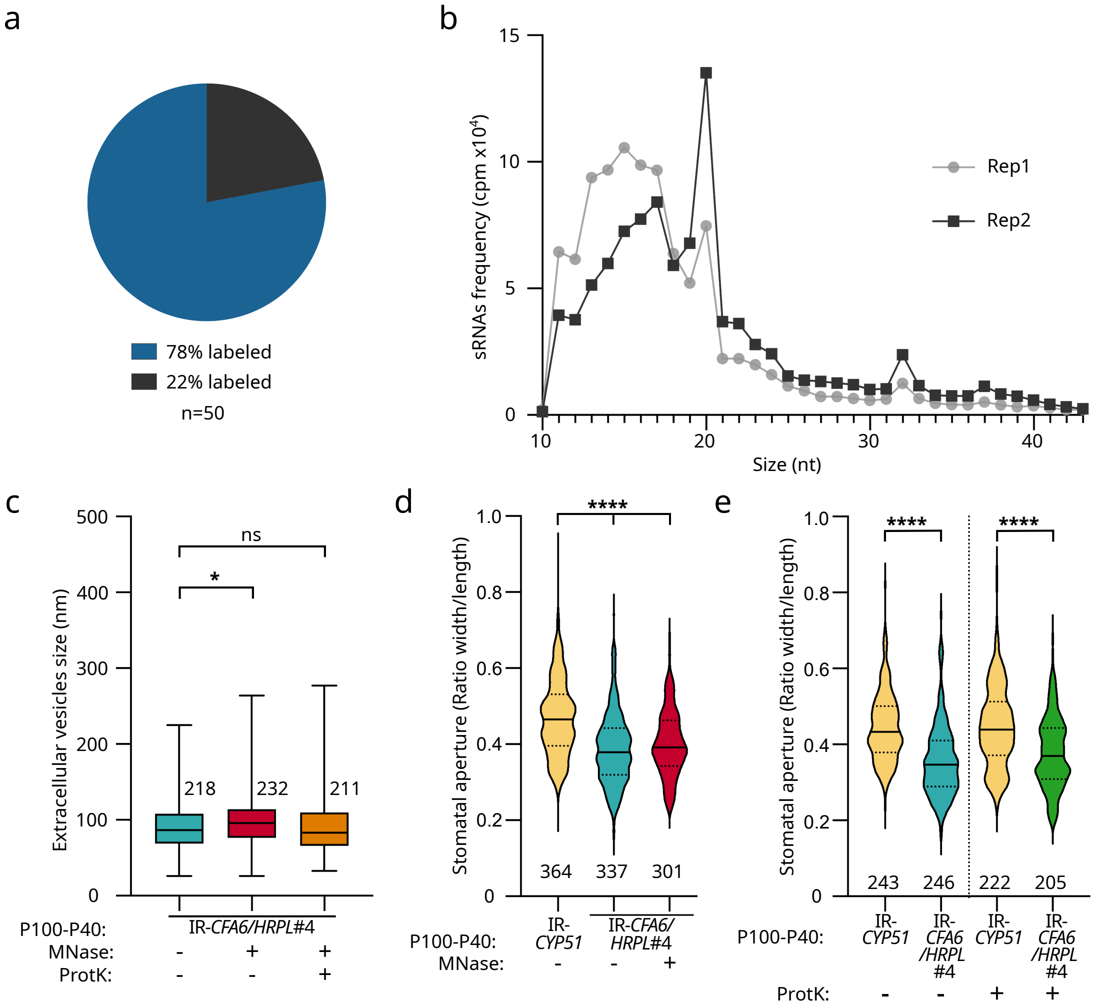

**Supplementary Figure 5. Characterization of the P100-P40 pellet, anti-*cfa6*/*hrpL* sRNAs accumulation and impact of MNase and Proteinase K treatments on TET8-positive EVs integrity.**

**a.** Percentage of EVs in the P100-P40 sample labeled with TET8 antibody. Immunolabeling was performed using TET8 antibodies and secondary antibodies coupled with gold beads. EVs were observed through TEM. **b.** Size distribution and abundance of sRNA reads from IR-*CFA6/HRPL*#4 MNase-treated P100-P40 pellets. Data from two biological replicates are presented, and the same replicates are used in Fig. 5b. **c.** Pictures of representative EVs were measured using ImageJ software. The number of vesicles measured is written above each condition and statistical significance was assessed using the ANOVA test (ns: p-value ≥ 0.05; *: 0.05 > p-value ≥ 0.01). Similar results were obtained in three independent experiments and the data are pooled in a single plot. **d.** MNase treatment alone does not affect P100-P40 pellet activity against *Pto* DC3000. Stomatal aperture measurements were conducted in Col-0 pre-treated with IR-*CFA6/HRPL*#4- or IR-*CYP51*-derived P100-P40 pellets that were subjected to enzymatic treatments. Leaf sections were infected with the *Pto* WT strain. The number of stomata analyzed per condition is written underneath each condition and statistical significance was assessed using a one-way ANOVA test (****: p-value < 0.0001). Similar results were obtained in three independent experiments and the data are pooled in a single plot. **e.** ProtK treatment alone does not affect P100-P40 pellet activity against *Pto* DC3000. The stomatal reopening assay was performed as described in d. Similar results were obtained in two independent experiments and the data are pooled in a single plot.

**Supplementary Figure 6**

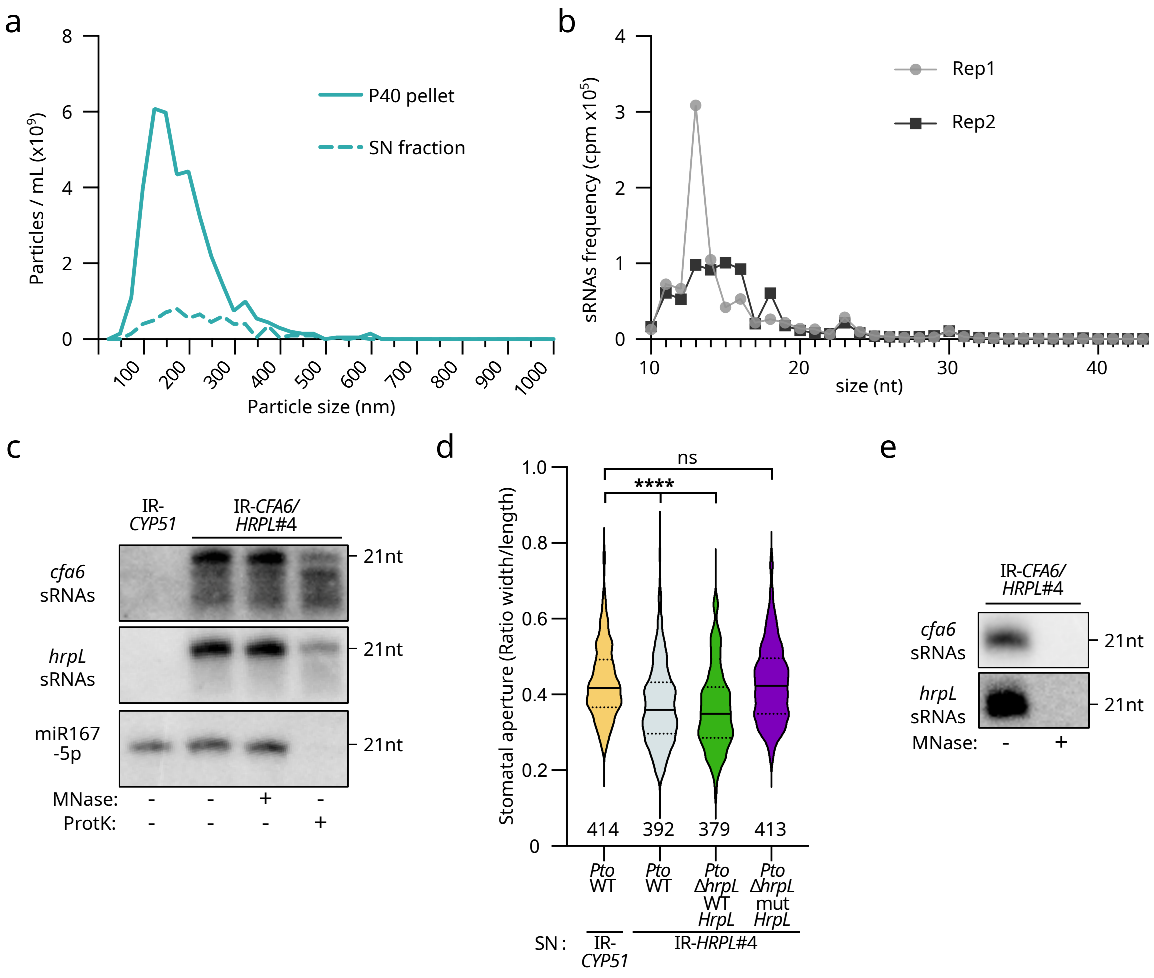

**Supplementary Figure 6. Characterization of the apoplastic SN fractions of IR-*CFA6*/*HRPL*#4 and IR-*CYP51* plants.**

**a.** Size distribution and abundance of nanoparticles in IR-*CFA6/HRPL*#4 P40 pellets or SN fraction. The analyses were performed using Nanoparticle Tracking Analysis (NTA) method. Similar results were obtained in three independent experiments. This is one representative experiment out of the three independent experiments performed. **b.** Size distribution and abundance of sRNA reads from the SN fraction of the IR-*CFA6/HRPL*#4 transgenic line. Data from two biological replicates are presented, and the same replicates are used in Fig. 6c. **c.** Anti-*cfa6* and anti-*hrpL* sRNAs accumulation is reduced after ProtK but not after MNase treatments, at the resolution of northern analysis. Accumulation of sRNAs in IR-*CFA6/HRPL*#4- and IR-*CYP51*-derived SN fractions was detected by low molecular weight northern blot analysis. MicroRNA167 (miR167-5p) was used as an endogenous miRNA control. This is one experiment out of the two independent experiments performed. **d.** The sRNAs from the SN fraction of IR-*HRPL*#4 plants suppress *Pto* DC3000-triggered stomatal aperture in a sequence-specific manner. Stomatal aperture measurements were conducted on Col-0 leaf sections incubated with IR-*HRPL*#4 or IR-*CYP51*-derived SN fractions. Leaf sections were inoculated with indicated bacterial strains and the stomatal aperture measurements were performed as in Supplementary Fig. 3e. Similar results were obtained in three independent experiments and are pooled in a single plot. **e.** MNase treatments fully degrade anti-*cfa6* and anti-*hrpL* sRNAs from total RNA extracts from IR-*CFA6/HRPL*#4 plants. Ten micrograms of IR-*CFA6/HRPL*#4 total RNAs were treated with MNase and anti-*cfa6/hrpL* sRNAs were further detected by low molecular weight northern blot analysis. This is one representative experiment out of the three independent experiments performed.

### Supplementary Figure 7

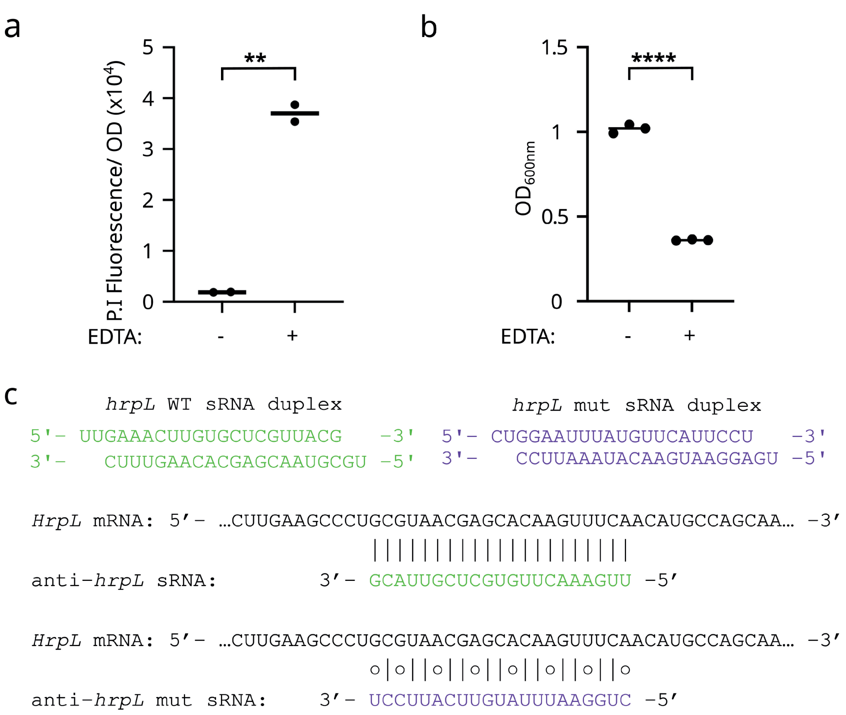

**Supplementary Figure 7. Effect of EDTA treatment on *Pto* DC3000, and internalized anti-*cfa6/hrpL* 21nt long sRNAs.**

**a.** Treatment with EDTA effectively degrades the outer membrane of *Pto* DC3000. Membrane permeability was evaluated by measuring the internalization of propidium iodide (P. I) using fluorescence. The fluorescence signal was normalized to the optical density at OD at 600 nm (OD). **b.** *Pto* DC3000 exhibits impaired growth after treatment with EDTA. The bacteria were treated with EDTA to disrupt the outer membrane and periplasm, then cultured in NYGB medium for 24 hours. *In vitro* growth was monitored by measuring $OD_{600}$. **c.** Upper panels: representations of the sRNA duplex with 2-nt 3' overhangs for the WT *hrpL* duplex (*hrpL* WT sRNA duplex) depicted in green, and its cognate mutated version (*hrpL* mut sRNA duplex), depicted in purple. Anti-*hrpL* WT sRNA (shown in green) is 100% complementary to *HrpL* mRNA, whereas anti-*hrpL* mut sRNA (shown in purple) has been mutated to alter anti-*hrpL* sRNA pairing with the *HrpL* mRNA.

**Supplementary Figure 8**

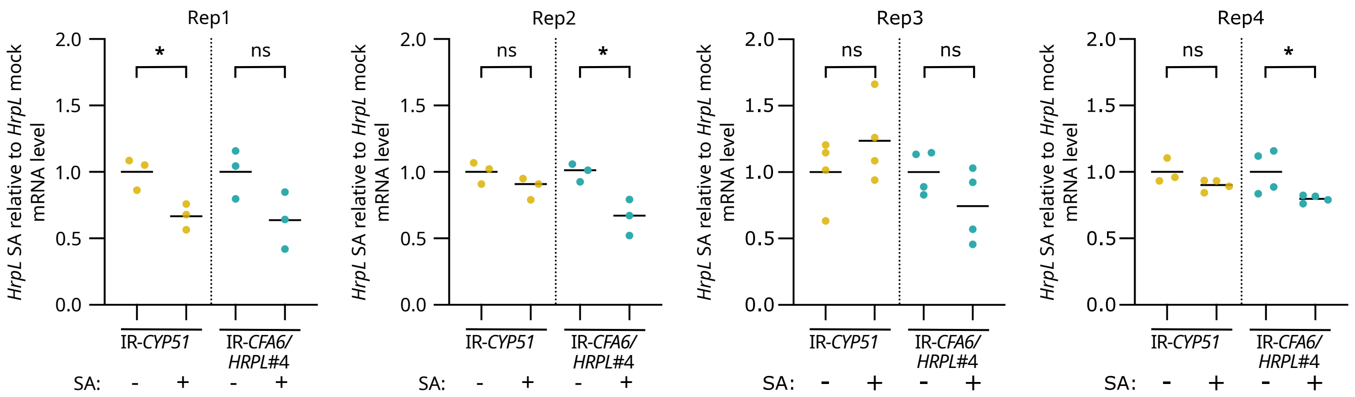

**Supplementary Figure 8. *HrpL* mRNAs are not consistently and significantly altered in *Pto* DC3000 cells in the presence of leaf sections from IR-*CFA6*/*HRPL*#4 and IR-*CYP51* plants pretreated with salicylic acid.**

RT-qPCR analyses depicting the levels of *HrpL* mRNAs in four independent experiments. The *HrpL* mRNA levels display slight or no significant changes in the presence of extra-cellular/organismal sRNAs from IR-*CFA6*/*HRPL*#4 leaf sections (with or without SA). Four independent biological replicates are represented. Levels of *HrpL* mRNAs in *Pto* DC3000 cells exposed to either IR-*CYP51* SA or IR-*CFA6*/*HRPL*#4 SA were normalized to IR-*CYP51* mock or IR-*CFA6*/*HRPL*#4 mock, respectively. Statistical significance was assessed using Student's t-test (ns: p-value ≥ 0.05; *: 0.05 > p-value ≥ 0.01).
